# Therapeutic Deep Brain Stimulation Disrupts Subthalamic Nucleus Activity Dynamics in Parkinsonian Mice

**DOI:** 10.1101/2021.11.12.468404

**Authors:** Jonathan S Schor, Isabelle Gonzalez Montalvo, Perry W E Spratt, Rea J Brakaj, Jasmine A Stansil, Kevin J Bender, Alexandra B Nelson

**Affiliations:** Neuroscience Program, University of California, San Francisco, San Francisco, CA 94158, USA; Department of Neurology, University of California, San Francisco, San Francisco, CA 94158, USA; Kavli Institute for Fundamental Neuroscience, University of California, San Francisco, San Francisco, CA 94158, USA; Weill Institute for Neuroscience, University of California, San Francisco, San Francisco, CA 94158, USA

## Abstract

Subthalamic nucleus deep brain stimulation (STN DBS) relieves many motor symptoms of Parkinson’s Disease (PD), but its underlying therapeutic mechanisms remain unclear. Since its advent, three major theories have been proposed: (1) DBS inhibits the STN and basal ganglia output; (2) DBS antidromically activates motor cortex; and (3) DBS disrupts firing dynamics within the STN. Previously, stimulation-related electrical artifacts limited mechanistic investigations using electrophysiology. We used electrical artifact-free calcium imaging to investigate activity in basal ganglia nuclei during STN DBS in parkinsonian mice. To test whether the observed changes in activity were sufficient to relieve motor symptoms, we then combined electrophysiological recording with targeted optical DBS protocols. Our findings suggest that STN DBS exerts its therapeutic effect through the disruption of STN dynamics, rather than inhibition or antidromic activation. These results provide insight into optimizing PD treatments and establish an approach for investigating DBS in other neuropsychiatric conditions.

## Introduction

The basal ganglia are a group of interconnected subcortical structures long believed to control movement through modulation of neuronal firing. This rate-based model posits that reductions in basal ganglia output (globus pallidus interna [GPi] and substantia nigra pars reticulata [SNr]) facilitate movement. According to this model, the loss of dopaminergic input to the basal ganglia in Parkinson’s disease (PD) leads to pathological increases in GPi/SNr firing and impaired movement. Dopamine replacement therapies, such as with dopamine agonists or the dopamine precursor levodopa, reduce GPi/SNr firing in both PD patients and animal models (Hutchinson et al., 1997; Levy et al., 2001; Lozano et al., 2000; Papa et al., 1999), suggesting they may act by restoring normal firing rates. Given these findings and the rate model, it is paradoxical that deep brain stimulation (DBS) of the subthalamic nucleus (STN), an intrinsic basal ganglia nucleus with predominantly excitatory projections to GPi/SNr, is one of the most effective treatments for PD (Hickey and Stacy, 2016).

Investigations into how STN DBS impacts STN, GPi, and SNr activity have yielded conflicting results, and thus, how STN DBS exerts its therapeutic effects remains unclear. However, three major theories have been suggested (Chiken and Nambu, 2014): (1) STN DBS *inhibits* STN activity, consistent with the rate model; (2) STN DBS bypasses basal ganglia output by antidromically *exciting* cortical neurons projecting to the STN; or (3) STN DBS *disrupts* movement-related dynamics in the STN. Supporting (1), focal STN lesions relieve motor symptoms of PD (Andy et al., 1963; Bergman et al., 1990), and some groups have observed inhibition of STN, GPi, or SNr firing during DBS (Filali et al., 2004; Moran et al., 2011). However, other groups have observed excitation in these structures (Hashimoto et al., 2003; Reese et al., 2011). Supporting (2), antidromic activation of primary motor cortex (M1) during STN DBS has been observed in rodents and in humans (Li et al., 2012; Miocinovic et al., 2018), and optical stimulation of hyperdirect M1 neurons relieves motor symptoms in parkinsonian mice (Gradinaru et al., 2009; Sanders and Jaeger, 2016). However, recent evidence in nonhuman primates shows antidromic M1 activation during electrical STN DBS to be both variable and transient, suggesting this pathway may not be the primary mechanism of STN DBS (Johnson et al., 2020). Supporting (3), correlations have been found between parkinsonian motor symptoms and signals such as local field potential (LFP) oscillations (Kühn et al., 2008; Stein and Bar-Gad, 2013; Wingeier et al., 2006) and bursting (Pan et al., 2016; Zhuang et al., 2018). Despite these observations, it has been difficult to distinguish between these possible mechanisms using electrophysiology, or to causally link observed changes in pattern and rhythm with improvements in behavior, in part due to large DBS-related electrical artifacts in recordings near the DBS site.

Here we used the genetically encoded calcium indicator GCaMP to enable region- and cell type-specific (and electrical artifact-free) optical recording of neural activity in three basal ganglia circuit nodes (STN, SNr, and M1) in a mouse model of STN DBS for PD. Surprisingly, we found that STN DBS *increased* activity in the STN and the SNr (in conflict with the rate model), and though DBS also altered hyperdirect pathway M1 activity, these changes did not correlate strongly with motor improvement. Furthermore, M1 lesions did not eliminate the therapeutic benefit of STN DBS. Finally, we found that DBS abolished stereotyped patterns of STN activity around movement onset. An optical stimulation protocol that similarly attenuated this activity pattern was sufficient to improve movement, suggesting disruption of movement-related STN dynamics may be a core therapeutic mechanism of STN DBS. Together, our results suggest STN DBS causes specific disruptions in motor signals at the level of the STN, broadening our understanding of how the basal ganglia mediates motor control.

## Results

### STN GCaMP signals correlate with spiking measured by electrophysiology

To test whether DBS inhibits, excites, or disrupts its target neurons, we used fiber photometry to measure bulk fluorescence signals from the genetically encoded calcium indicator GCaMP6s. To validate this approach, we first sought to determine whether STN calcium signals could serve as proxy for neural activity. We injected VGlut2-Cre mice with AAVs encoding Cre-dependent GCaMP6s, limiting expression to glutamatergic neurons within the STN. We then performed simultaneous whole-cell current-clamp recordings and fluorescence imaging of STN neurons in *ex vivo* slices to compare firing rate and intracellular calcium signals (Fig. 1A-C). Neurons were stimulated with current pulses at frequencies ranging from 10 Hz to 120 Hz in 1-minute epochs, which evoked rhythmic spiking corresponding to the frequency of stimulation (Fig. 1D-E). Calcium, as measured by changes in GCaMP6s fluorescence, similarly increased during stimulation at 10-120 Hz, though notably in a non-linear fashion, suggesting a weaker correspondence between spiking and calcium at the highest stimulation frequencies (Fig. 1F-G). This phenomenon may also be related to limitations in GCaMP6s signaling at high intracellular calcium levels. In a subset of recordings, we assessed GCaMP6s signals in response to a range of lower stimulation frequencies (10-60 Hz). Again, STN neurons showed rhythmic spiking that matched the frequency of pulsatile stimulation (Fig. 1E, inset), and calcium signals correlated with firing rates (Fig. 1G, inset). In response to constant current (“square wave”) stimulation, STN neurons fired only transiently, appearing to enter depolarization block (Fig. 1D-E). Under these circumstances, evoked calcium signals fell between those evoked by 10 and 50-60 Hz stimulation (Fig. 1F-G). These experiments suggest that the relationship between spiking and GCaMP calcium signals may break down at very high frequencies of stimulation, or under conditions of forced depolarization block. However, at the more moderate frequencies explored here, GCaMP calcium signals correlate with STN firing.

**Figure 1.**
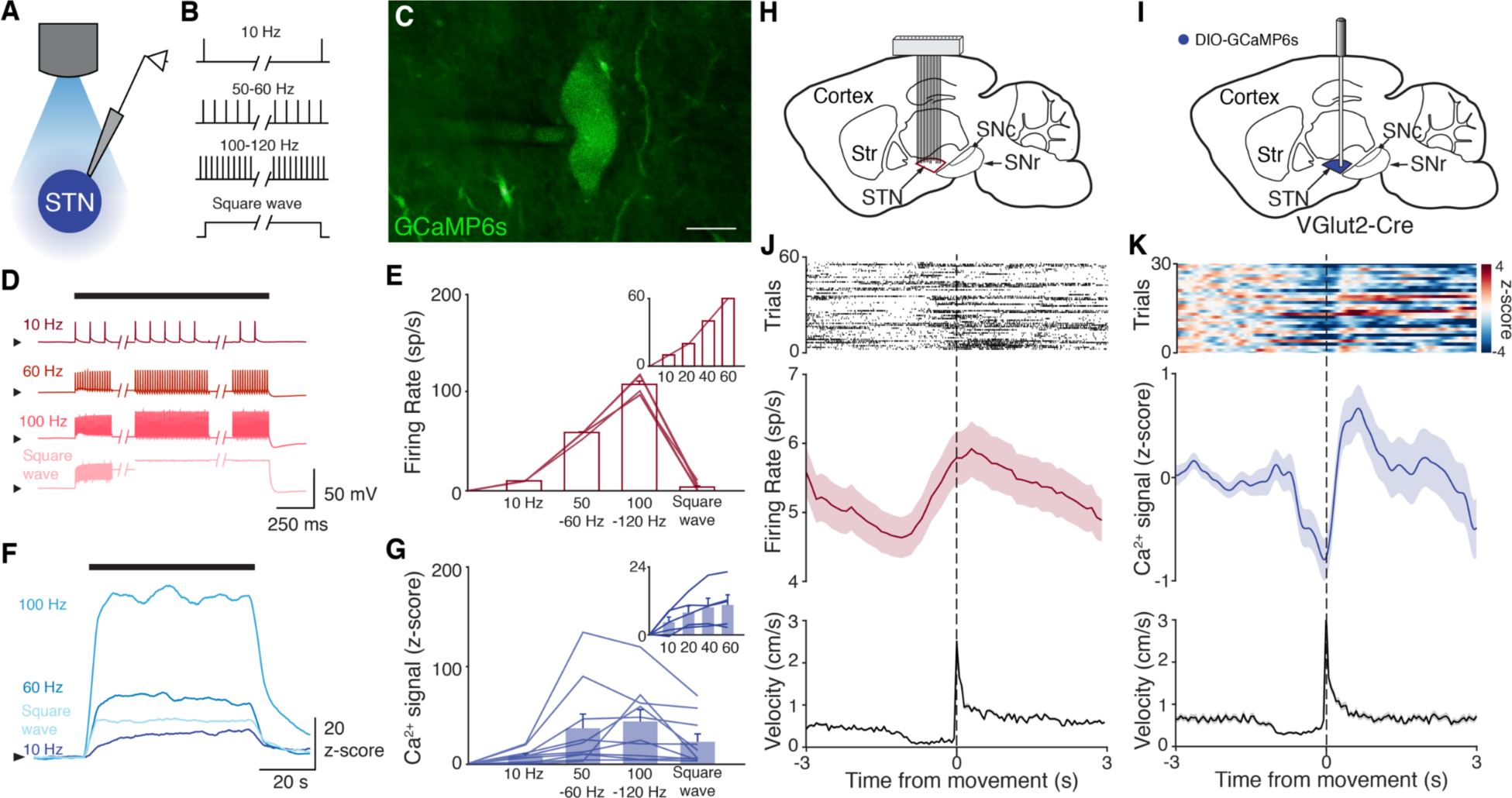
STN GCaMP signals correlate with spiking measured by electrophysiology. (A-G) Combined ex vivo electrophysiological and calcium imaging recordings in STN neurons. Neurons were patched in the whole-cell current-clamp configuration. **(A)** Recording configuration. **(B)** Schematic showing current-clamp stimulation protocol. **(C)** Image of GCaMP-expressing STN neuron (scale bar=10 μm). **(D)** Representative STN neuron responses to the indicated current-clamp stimulation. **(E)** Average firing rate of STN neurons in response to stimulation (n=10 cells, N=3 mice for main; n=5 cells, N=1 mouse for inset). **(F)** Representative trace of z-scored STN GCaMP signal in response to current-clamp stimulation. **(G)** Average z-scored STN GCaMP signal in response to current-clamp stimulation (n=10 cells, N=3 mice for main; n=5 cells, N=1 mouse for inset). Arrowhead in current-clamp traces and GCaMP traces corresponds to -75 mV and 0 z-score, respectively. Bar plots show mean ± SEM. (H-K) *In vivo* electrophysiological and calcium imaging recordings in STN neurons from freely moving parkinsonian mice, aligned to movement starts. **(H)** Sagittal schematic showing multielectrode array implant in the STN for single-unit electrophysiology. **(I)** Sagittal schematic showing STN GCaMP and fiber implant for photometry. **(J)** Representative STN single-unit firing (top), average firing rate (middle), and average velocity (bottom) aligned to movement starts (n=17 cells, N=3 mice). **(K)** Representative STN fiber photometry signal (top), average fiber photometry signal (middle), and average velocity (bottom) aligned to movement starts (N=8 mice). Average firing rate, photometry, and velocity traces show mean ± SEM.

We next tested whether changes in bulk GCaMP6s fluorescence, as recorded through *in vivo* fiber photometry, corresponded with *in vivo* single-unit activity. To address this question, we recorded neural activity *in vivo* in two sets of parkinsonian animals, using either electrophysiology or calcium imaging. We rendered mice parkinsonian through unilateral injection of 6-OHDA in the medial forebrain bundle (MFB, Supplementary Fig. 1A). In one group of parkinsonian mice, we implanted a 16-channel electrode array in the ipsilateral STN (Fig. 1H). In a second group of parkinsonian VGlut2-Cre mice, we injected Cre-dependent GCaMP6s and implanted an optical fiber in the ipsilateral STN (Fig. 1I). As observed in prior 6-OHDA studies (Bové and Perier, 2012; Campos et al., 2013; Carvalho et al., 2013; Ungerstedt, 1968), these mice showed both decreased movement velocity (1.3 ± 0.1 cm/s parkinsonian vs 3.1 ± 0.4 cm/s healthy, *P*=8.23x10^-5^) and an ipsilesional rotational bias (*P*=1.40x10^-3^) when compared to healthy mice (Supplementary Fig. 1B). Additionally, movement velocity and rotation bias did not differ significantly between parkinsonian mice with and without STN implants (Supplementary Fig. 1B, *P*=0.66 for velocity, *P*=0.97 for rotation), suggesting local, implant-related STN tissue disturbance did not alter gross movement parameters. We then aligned single-unit spiking activity and fiber photometry signal to movement starts (defined as a transition from velocity <0.5 cm/s to >2 cm/s) in both sets of mice (Fig. 1J-K). STN single units showed baseline firing rates of approximately 5-10 spikes/s (Fig. 1J), well within the linear range for GCaMP6s (Fig. 1G, inset). The firing of STN units showed a marked change in firing rate around movement onset, increasing just prior to, and peaking just following movement initiation (approximately 1 spike/s over the baseline rate). Population calcium signals showed a similar increase around movement onset, but lagged the rise in single-unit firing rate by ∼1 sec, likely due to the slower kinetics of GCaMP6s (Markowitz et al., 2018) (Fig. 1K). Thus, STN calcium signals and electrophysiology appear to capture a slightly time-shifted, but qualitatively similar increase in activity when aligned to behavior. Together, these *ex vivo* and *in vivo* experiments show a correlation between spiking and GCaMP6s signal, and therefore support the utility of fiber photometry as a proxy for neuronal activity in the context of STN DBS.

### STN DBS consistently increases STN activity

The direct impact of STN DBS on STN neural activity remains unclear: some studies indicate STN DBS decreases STN firing rates, while recordings in downstream nuclei indicate STN activity may increase. To address whether STN DBS increases or decreases overall STN activity, we injected parkinsonian VGlut2-Cre mice with Cre-dependent GCaMP6s and implanted them with both an STN DBS device and an optical fiber (Fig. 2A-B, Supplementary Fig. 2A,3A). Consistent with our previous findings in the same mouse model (Schor and Nelson, 2019), electrical STN DBS improved multiple movement metrics (Supplementary Fig. 2B-K). At either 60 or 100 Hz stimulation, STN DBS increased movement velocity (Supplementary Fig. 2B,C,G,H; *P*=3.92x10^-9^ for 60 Hz, *P*=5.53x10^-8^ for 100 Hz) and percent time moving (Supplementary Fig. 2D,I; *P*=3.53x10^-9^ for 60 Hz, *P*=1.46x10^-8^ for 100 Hz), while not significantly altering rotation bias (Supplementary Fig. 2E,J; *P*=0.71 for 60 Hz, *P*=0.47 for 100 Hz) or causing prolonged dyskinesias (Supplementary Fig. 2F,K). We subsequently used movement velocity as the primary behavioral outcome measure for STN DBS. We then measured changes in STN activity *in vivo* in response to treatment with STN DBS. Surprisingly, in parallel with its impact on movement velocity (*P*=7.11x10^-9^), STN DBS at 60 Hz caused a significant increase in STN calcium signals (Fig. 2C-D, Supplementary Fig. 3B; *P*=1.19x10^-7^). The same was true for stimulation at 100 Hz, with an increase in STN activity (*P*=9.54x10^-7^) mirroring an increase in velocity (Fig. 2E-F, Supplementary Fig. 3C; *P*=1.11x10^-9^). This result suggests that, rather than inhibiting the STN, STN DBS increases STN activity.

**Figure 2.**
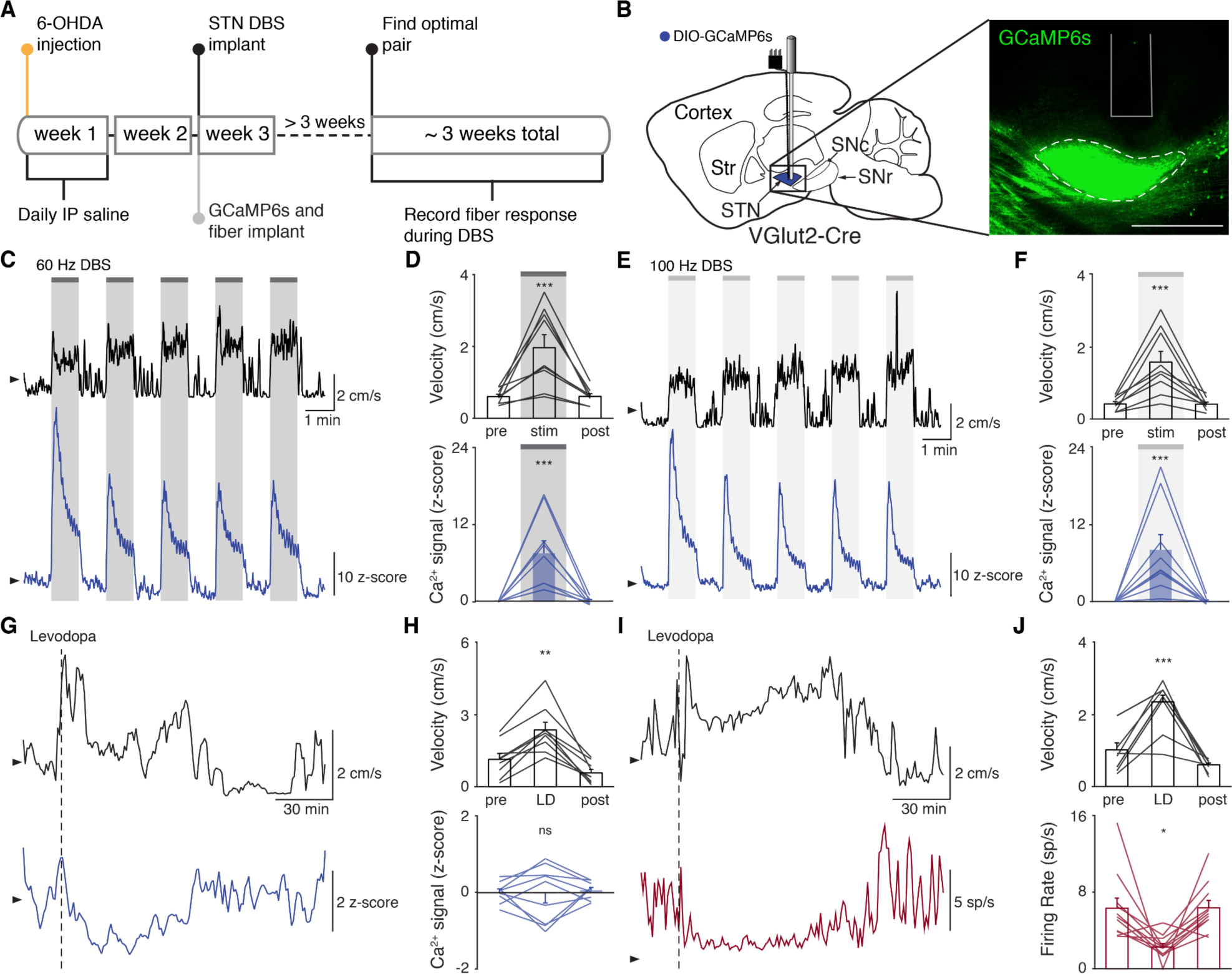
STN DBS consistently increases STN activity. **(A)** Experimental timeline. **(B)** Left: Sagittal schematic showing STN DBS and GCaMP fiber photometry. Right: Postmortem sagittal section showing GCaMP expression and estimated fiber placement in the STN (inset, scale=500 μm). **(C)** Representative single-session velocity (black) and STN GCaMP signal (blue) in response to 60 Hz STN DBS. **(D)** Average velocity (top) and STN GCaMP signal (bottom) before, during, and after 60 Hz STN DBS (N=9 mice). **(E)** Representative single-session velocity (black) and STN GCaMP signal (blue) in response to 100 Hz STN DBS. **(F)** Average velocity (top) and STN GCaMP signal (bottom) before, during, and after 100 Hz STN DBS (N=9 mice). **(G)** Representative single-session velocity (black) and STN GCaMP signal (blue) before and after levodopa injection (dotted line). **(H)** Average velocity (top) and STN GCaMP signal (bottom) before, during, and after levodopa treatment (N=9 mice). **(I)** Representative single-session velocity (black) and STN single-unit activity (red) before and after levodopa injection (dotted line). **(J)** Average velocity (top) and STN single-unit activity (bottom) before, during, and after levodopa treatment (n=11 cells, N=3 mice). Statistical significance was determined using a one-way repeated measures ANOVA with a Tukey HSD post hoc analysis applied to correct for multiple comparisons; \**P* < 0.05, \*\**P* < 0.01, \*\*\**P* < 0.001 (only comparison between pre and stim/LD shown, see Supplementary Table 1 for detailed statistics). Arrowhead in velocity, GCaMP, and single-unit electrophysiology traces corresponds to 1 cm/s, 0 z-score, and 0 sp/s, respectively. Bar plots show mean ± SEM.

These results appear to conflict with the proposed inhibitory mechanism of other Parkinson’s disease treatments, such as surgical ablations and dopamine replacement therapy. To compare how STN DBS and dopamine replacement therapy impact STN activity, we next evaluated the effects of levodopa administration on STN activity. In the same set of mice used for testing STN DBS, levodopa increased movement velocity (Supplementary Fig. 2L-M; *P*=9.69x10^-10^) and percent time moving (Supplementary Fig. 2N; *P*=9.56x10^-10^), evoked contralesional rotations (Supplementary Fig. 2O; *P*=1.73x10^-6^), and caused minimal dyskinesias (Supplementary Fig. 2P). We subsequently used movement velocity and rotation bias as primary and secondary behavioral outcome measures of levodopa treatment, respectively. In these sessions, though all mice showed improvements in movement (*P*=1.66x10^-3^ for velocity, *P*=3.91x10^-3^ for rotation bias), changes in STN calcium were variable (Fig. 2G-H; Supplementary Fig. 3D). In some mice, STN activity decreased, while in others it increased: STN activity was not significantly changed across the entire group (Fig. 2H; *P*=0.90). Injection with saline did not improve movement parameters, nor did it produce significant changes in STN activity (Supplementary Fig. 3E-F; *P*=0.084). As levodopa does not produce electrical artifacts like DBS, we were able to compare its effects on both calcium signals and single-unit firing rates using electrophysiology. In parallel with its impact on movement (*P*=3.13x10^-4^ for velocity, *P*=9.77x10^-4^ for rotation bias), levodopa caused a modest, though significant, decrease in STN firing rates (Fig. 2I-J, Supplementary Fig. 3G; *P*=0.029). The fact that bulk calcium imaging did not detect the modest reductions in STN firing seen with electrophysiology may relate to differential sensitivity of the two methods. However, despite the fact that DBS and levodopa both improve movement parameters, they alter overall activity level in opposite directions.

### STN DBS increases SNr activity

Though the STN is a critical node within the basal ganglia circuit, especially in regard to dysfunction in PD and its treatment, changes in STN activity are believed to regulate motor function via excitatory projections to basal ganglia output nuclei. In addition, STN DBS is likely to cause changes in the activity of nearby axons, and thus may have complex downstream effects. Therefore, while we did not observe inhibition at the level of the STN during STN DBS, we wondered if it might still produce inhibition at the level of the primary basal ganglia output nucleus in rodents, the substantia nigra pars reticulata (SNr). To determine how SNr activity responds to STN DBS, we injected either VGAT-Cre mice with Cre-dependent GCaMP6s in the SNr (N=6 mice) or WT mice with synapsin-GCaMP6s in the SNr (N=2 mice) and implanted them with an STN DBS device and an optical fiber in the SNr (Fig. 3A, Supplementary Fig. 4A). As before, STN DBS in these mice increased movement velocity (Fig. 3B-E; *P*=1.33x10^-5^ for 60 Hz, *P*=4.99x10^-6^ for 100 Hz). Consistent with our results in the STN, STN DBS (at both 60 and 100 Hz) increased SNr activity (Fig. 3B-E; Supplementary Fig. 4B-C; *P*=4.14x10^-5^ for 60 Hz, *P*=3.32x10^-5^ for 100 Hz). Contrary to the basal ganglia rate model, and the inhibition theory of DBS, these findings suggest that both STN and SNr activity are increased by STN DBS.

**Figure 3.**
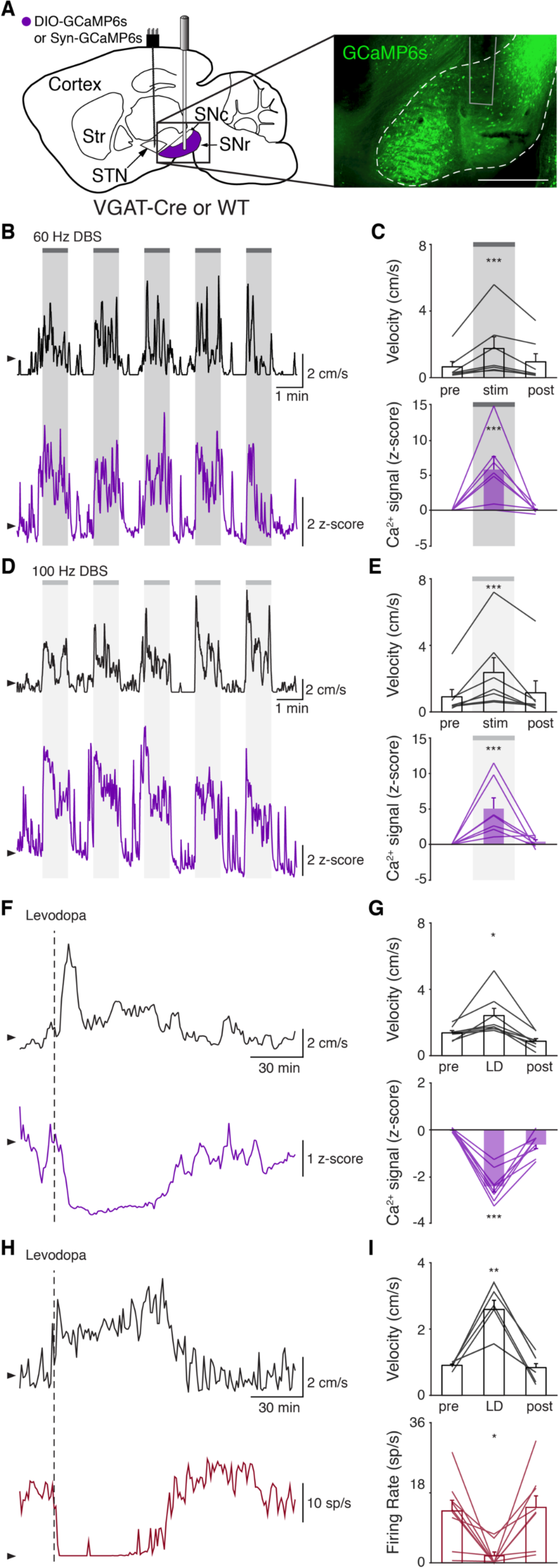
STN DBS increases basal ganglia output. **(A)** Left: Sagittal schematic showing STN DBS and SNr GCaMP fiber photometry. Right: Postmortem sagittal section showing GCaMP expression and estimated fiber placement in the SNr (inset, scale=500 μm). **(B)** Representative single-session velocity (black) and SNr GCaMP signal (purple) in response to 60 Hz STN DBS. **(C)** Average velocity (top) and SNr GCaMP signal (bottom) before, during, and after 60 Hz STN DBS (N=7 mice). **(D)** Representative single-session velocity (black) and SNr GCaMP signal (purple) in response to 100 Hz STN DBS. **(E)** Average velocity (top) and SNr GCaMP signal (bottom) before, during, and after 100 Hz STN DBS (N=7 mice). **(F)** Representative single-session velocity (black) and SNr GCaMP signal (purple) before and after levodopa injection (dotted line). **(G)** Average velocity (top) and SNr GCaMP signal (bottom) before, during, and after levodopa treatment (N=8 mice). **(H)** Representative single-session velocity (black) and SNr single-unit activity (red) before and after levodopa injection (dotted line). **(I)** Average velocity (top) and SNr single-unit activity (bottom) before, during, and after levodopa treatment (n=9 cells, N=3 mice). Statistical significance was determined using a one-way repeated measures ANOVA with a Tukey HSD post hoc analysis applied to correct for multiple comparisons; \**P* < 0.05, \*\**P* < 0.01, \*\*\**P* < 0.001 (only comparison between pre and stim/LD shown, see Supplementary Table 1 for detailed statistics). Arrowhead in velocity, GCaMP, and single-unit electrophysiology traces corresponds to 1 cm/s, 0 z-score, and 0 sp/s, respectively. Bar plots show mean ± SEM.

To again validate calcium imaging signals and compare DBS to other treatments, we measured how levodopa altered SNr activity. In the same parkinsonian mice, levodopa increased movement velocity (*P*=0.016) and caused a contralesional rotation bias (Fig. 3F-G; Supplementary Fig. 4D; *P*=7.81x10^-3^). In parallel, we observed a marked decrease in SNr neural activity as measured by calcium imaging (Fig. 3F-G; Supplementary Fig. 4D; *P*=6.68x10^-6^). In contrast, saline neither improved movement parameters nor significantly changed SNr calcium signals (Supplementary Fig. 4E-F; *P*=0.31). Single-unit electrophysiological recordings also showed profound reductions in SNr firing rate (*P*=0.012) during therapeutic levodopa treatment (Fig. 3H-I; Supplementary Fig. 4G), similar to findings in the GPi of parkinsonian nonhuman primates (Papa et al, 1999). Thus, calcium imaging and electrophysiology revealed qualitatively similar changes in SNr neural activity in response to levodopa, again supporting the idea that these two measures of neural activity have substantial concordance. Furthermore, these imaging experiments show marked differences in how STN DBS and levodopa impact neural activity, suggesting that STN DBS does not exert therapeutic effects through inhibition of STN or SNr.

### Hyperdirect Pathway Activity During STN DBS

While we did not observe inhibition in either the STN or SNr during STN DBS, a more recent theory posits that STN DBS acts through antidromic activation of the hyperdirect pathway: primary motor cortex (M1) neurons that project monosynaptically to the STN. To assess whether STN DBS increases activity of hyperdirect M1 neurons, we imaged hyperdirect pathway neurons using a retrograde viral strategy. We injected the STN of parkinsonian mice with one of two retrograde viruses encoding Cre recombinase (CAV2-Cre or rAAV2-Cre-mCherry), and injected M1 with Cre-dependent GCaMP6s (Fig. 4A, Supplementary Fig. 5A-B). This strategy restricted expression of GCaMP6s to STN-projecting M1 neurons, which previously have been shown to send collaterals to the STN with parent axons in the cerebral peduncle (Kita and Kita, 2012) (Supplementary Fig. 5B). We then implanted an optical fiber in M1 and a DBS device in the STN. As in other parkinsonian mice, both 60 and 100 Hz STN DBS increased movement velocity (Fig. 4B-E; *P*=1.51x10^-9^ for 60 Hz, *P*=4.14x10^-7^ for 100 Hz). Despite the consistent therapeutic effects of 60 Hz STN DBS (Fig. 4C, top), hyperdirect M1 calcium responses were surprisingly variable: some mice showed increases, while the calcium signal in other mice decreased or did not change (Fig. 4C, bottom; Supplementary Fig. 5C; *P*=0.084). In the same mice, 100 Hz STN DBS also produced consistent increases in movement velocity (Fig. 4E, top), but in this case M1 activity was more consistently increased during stimulation (Fig. 4E, bottom; Supplementary Fig. 5D; *P*=5.88x10^-5^). These results suggested poor correlation between hyperdirect M1 activity and the behavioral benefits of STN DBS. To further probe the correspondence between DBS effectiveness and hyperdirect M1 activation, we asked if the movement velocity of a single mouse during 60 Hz STN DBS could be predicted by that mouse’s M1 calcium activity. We found that movement velocity during DBS did not correlate with change in hyperdirect M1 neural activity (Supplementary Fig. 5E; R^2^=-0.14, *P*=0.96). These findings suggest that while certain stimulation parameters may promote hyperdirect pathway activity, these changes do not strongly correlate with behavioral improvements during DBS.

**Figure 4.**
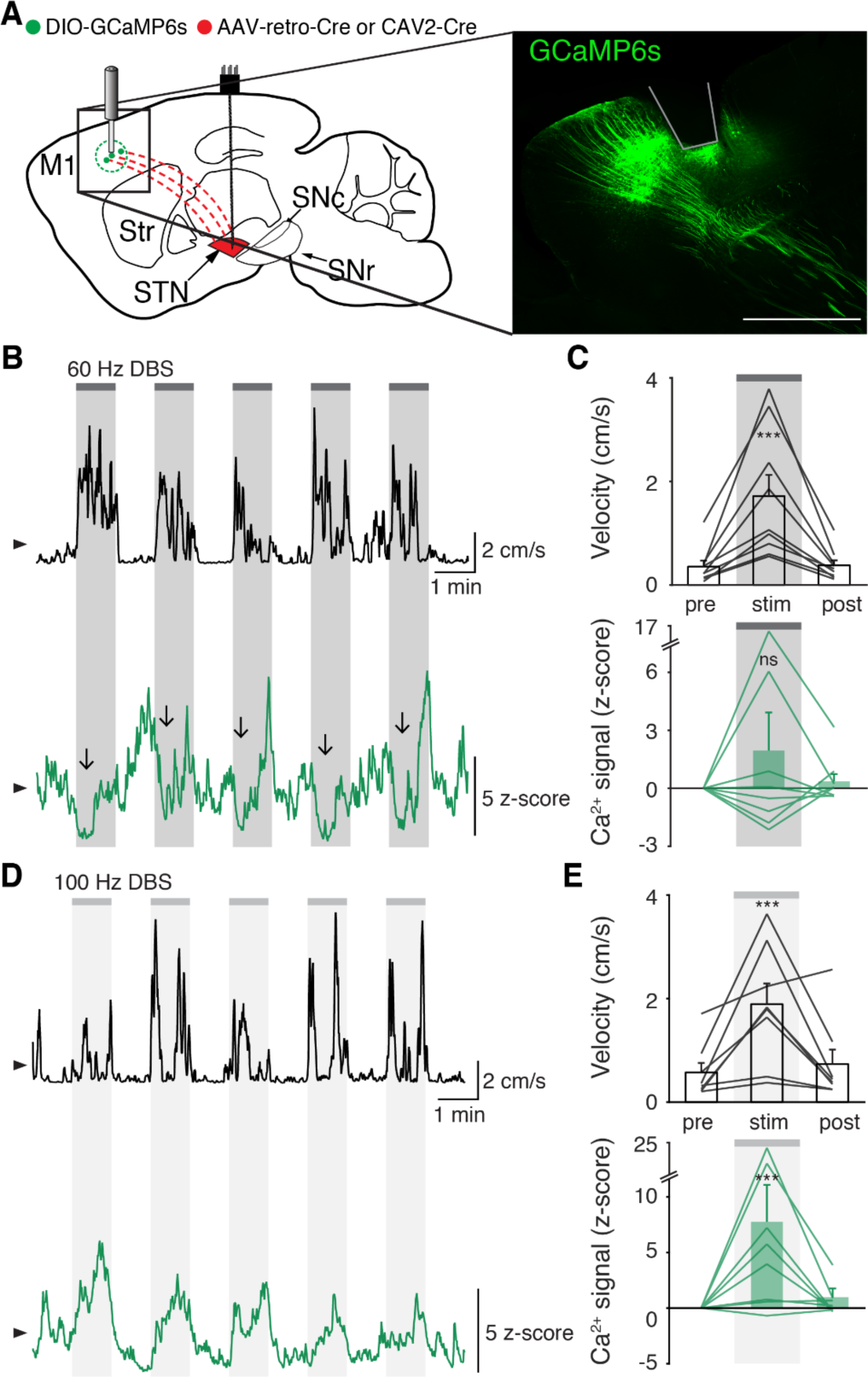
STN DBS variably changes hyperdirect M1 neural activity. **(A)** Left: Sagittal schematic showing STN DBS and M1-STN GCaMP fiber photometry. Right: Postmortem sagittal section showing GCaMP expression and estimated fiber placement in M1 (inset, scale=500 μm). **(B)** Representative single-session velocity (black) and M1-STN GCaMP signal (green) in response to 60 Hz STN DBS. **(C)** Average velocity (top) and M1-STN GCaMP signal (bottom) before, during, and after 60 Hz STN DBS (N=9 mice). **(D)** Representative single-session velocity (black) and M1-STN GCaMP signal (green) in response to 100 Hz STN DBS. **(E)** Average velocity (top) and M1-STN GCaMP signal (bottom) before, during, and after 100 Hz STN DBS (N=8 mice). Statistical significance was determined using a one-way repeated measures ANOVA with a Tukey HSD post hoc analysis applied to correct for multiple comparisons; \*\*\**P* < 0.001 (only comparison between pre and stim shown, see Supplementary Table 1 for detailed statistics). Arrowhead in velocity and GCaMP traces corresponds to 1 cm/s and 0 z-score, respectively. Bar plots show mean ± SEM.

Additionally, we noted that increases in hyperdirect M1 activity evolved more slowly during STN DBS than increases in STN and SNr activity. The activity of neurons mediating the therapeutic effects of STN DBS would be predicted to change on a similar timescale to behavior. To compare the activation kinetics of STN, SNr, and hyperdirect M1 neurons during DBS, we measured the rise time of the calcium signal in regions and conditions in which we observed significant changes in neural activity: STN (60 and 100 Hz), SNr (60 and 100 Hz), and hyperdirect M1 (100 Hz). Given the observed lag between electrophysiology and bulk GCaMP signals seen by other groups (Markowitz et al., 2018), and in our own data (Fig. 1), we expected that changes in neural activity driving motor benefits (as measured with GCaMP) might appear to lag the behavior itself. Across all STN DBS conditions, the rise time for movement velocity averaged 2.8 ± 0.6 sec. For each condition, we calculated the difference in rise time between the calcium signal and movement velocity as an indicator of whether these two signals changed on a similar timescale. STN calcium signals during 60 Hz or 100 Hz STN DBS lagged movement velocity by 2.6 ± 1.1 sec. We observed a similarly short lag comparing SNr calcium signals to the corresponding movement velocity traces (3.7 ± 1.8 sec). However, the lag in hyperdirect M1 activity was markedly longer (17.3 ± 3.1 sec). These kinetics indicate STN and SNr activity evolve on a timescale similar to the movement benefits of STN DBS, while hyperdirect M1 activity, as measured by fiber photometry, evolves more slowly.

### Surgical removal of M1 does not abolish therapeutic benefit of STN DBS

Though overall hyperdirect pathway activity was not a strong predictor of the therapeutic effects of STN DBS, these findings do not exclude the possibility that the hyperdirect pathway mediates motor benefits. We next asked if M1 was required for the therapeutic effects of STN DBS on movement. We surgically removed the ipsilesional M1 of parkinsonian mice and implanted STN DBS devices (Fig. 5A, Supplementary Fig. 5F). As in previous motor cortex lesion studies (Kawai et al., 2015), mice were allowed to recover for at least 10 days before behavioral testing. Remarkably, these mice still showed a significant increase in movement velocity in response to 100 Hz STN DBS (Fig. 5B-C; *P*=8.24x10^-3^). Thus, it is unlikely that antidromic activation of M1 is the primary driver of the therapeutic benefit of STN DBS in parkinsonian mice.

**Figure 5.**
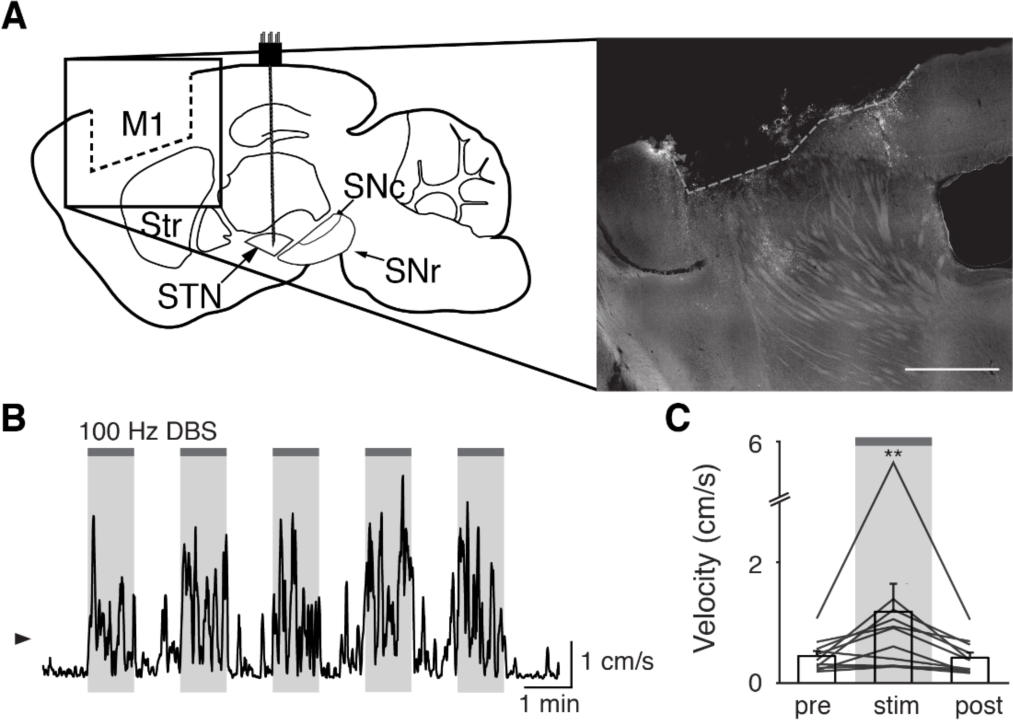
Surgical removal of M1 does not abolish therapeutic benefit of STN DBS. **(A)** Left: Sagittal schematic showing STN DBS and M1 surgical lesion. Right: Postmortem sagittal section showing estimated extent of M1 lesion (inset, scale=500 μm). **(B)** Representative single-session velocity of an M1-lesioned hemiparkinsonian mouse in response to 100 Hz STN DBS. **(C)** Average velocity before, during, and after 100 Hz STN DBS in M1-lesioned hemiparkinsonian mice (N=11 mice). Statistical significance was determined using a one-way repeated measures ANOVA with a Tukey HSD post hoc analysis applied to correct for multiple comparisons; \*\**P* < 0.01 (only comparison between pre and stim shown, see Supplementary Table 1 for detailed statistics). Arrowhead in velocity trace corresponds to 1 cm/s. Bar plots show mean ± SEM.

### STN movement-related activity is disrupted by therapeutic STN DBS

Our results indicate that STN DBS is unlikely to work through inhibition of basal ganglia output, nor through antidromic excitation of M1. However, a third possibility is that STN DBS disrupts neural dynamics within the STN itself. As previously noted, in parkinsonian mice, STN activity increases around movement initiation (Fig. 6A, left), consistent with the idea that STN neurons encode some aspects of movement. To determine whether this encoding was altered during DBS, we aligned neural activity to movement starts during stimulation epochs. We found that although the overall average calcium signal was increased during DBS (Fig. 2), the movement-aligned increase in STN activity was abolished during therapeutic STN DBS at 100 Hz (Fig. 6A, right). This observation suggested that STN movement encoding was disrupted by STN DBS.

**Figure 6.**
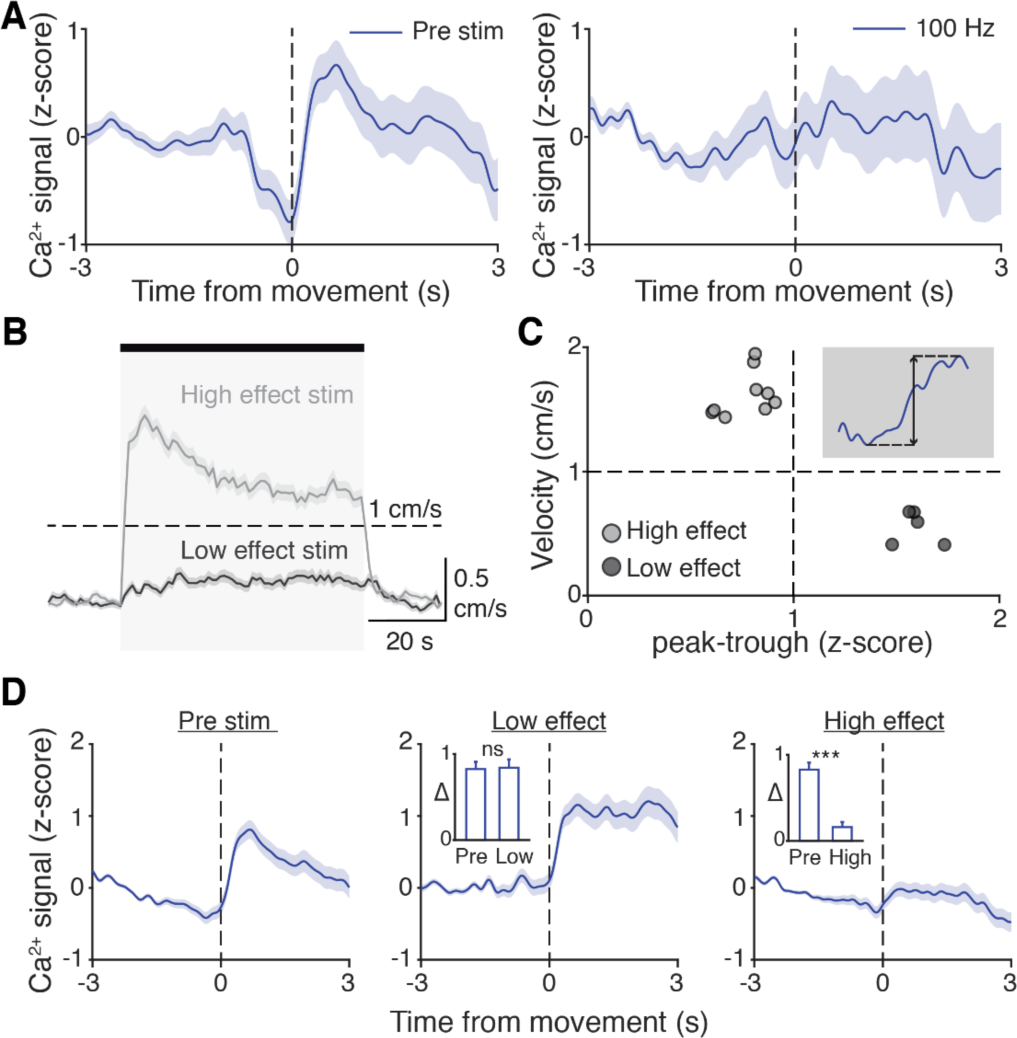
STN movement-related activity is disrupted by therapeutic STN DBS. **(A)** Average STN fiber photometry signal aligned to movement starts during pre-stim periods (left) and during 100 Hz STN DBS stimulation (right). **(B)** Average movement velocity over time during high effect stimulation (light grey, achieves average velocity >1 cm/s) and low effect stimulation (dark grey, achieves average velocity <1 cm/s). **(C)** Scatter plot comparing peak-trough distance in the average movement-aligned photometry trace (inset) to velocity during high effect (light grey) and low effect (dark grey) stimulation parameters. Each dot represents an average across all mice for each stimulation parameter (N=8 mice). **(D)** Average STN fiber photometry signal aligned to movement starts during pre-stim periods (left) and during low effect STN DBS stimulation (middle) and high effect STN DBS stimulation (right). Inset bar graphs show difference between average fiber signal in 1 sec following movement start and 1 sec preceding movement start (see Methods for further details) during pre, low effect, or high effect stim. Statistical significance was determined using a Wilcoxon rank-sum test; \*\*\**P* < 0.001 (see Supplementary Table 1 for detailed statistics). Bar plots, photometry, and velocity traces show mean ± SEM.

This disruption might result from any patterned electrical stimulation, or it could represent a direct correlate of therapeutic stimulation. To test these possibilities, we chose 13 parameter combinations (varying in current amplitude, frequency, and pulse width, Supplementary Table 2) from a set that we had used previously to evaluate the impact of STN DBS parameters on behavioral benefit in the mouse model (Schor and Nelson, 2019). In the same mice, we measured locomotor activity and calcium signals while delivering STN DBS at each of these 13 parameter combinations. Behavioral responses were divided into two groups, depending on whether the DBS-induced movement velocity averaged greater or less than 1 cm/s (Fig. 6B). We labeled the first group “high effect” stimulation (8 combinations) and the second group “low effect” stimulation (5 combinations). We then assessed STN movement-related activity dynamics during each stimulation type, as measured by the peak-to-trough deflection of the movement-aligned STN photometry signal (Fig. 6C, shaded inset). Interestingly, across the 13 stimulation parameter sets, this neural activity metric showed a bimodal distribution. With low effect stimulation parameters, STN calcium signals increased around movement onset, as they did during baseline (no stimulation) periods. This resulted in a peak-to-trough change in calcium that was >1 (z-scored dF/F; Fig. 6C). However, during high effect stimulation, STN calcium signals changed minimally around movement onset, with a peak-to-trough change of <1 (Fig. 6C). When comparing STN activity across all high effect vs all low effect stimulation parameters rather than individually, normal movement-related STN dynamics were strongly suppressed only during highly effective stimulation (Fig. 6D; *P*=1.48x10^-11^ for pre vs. high, *P*=0.54 for pre vs low). Taken together, these results suggest that STN DBS disrupts movement-related STN dynamics and furthermore that this disruption is specific to behaviorally beneficial stimulation parameters.

### Disruption of STN motor dynamics is sufficient to provide therapeutic benefit

STN DBS may trigger many changes in both the rate and dynamics of neural activity. However, it remains critical to determine which changes in neural activity causally contribute to the therapeutic mechanism(s) of STN DBS. In our experiments using electrical STN DBS, we observed changes in both overall STN activity and movement-related dynamics, making it difficult to determine which change is more likely to mediate the benefit. To disentangle the behavioral impacts of changing STN rate and dynamics, we replaced electrical DBS/optical recording with optical DBS/electrical recording techniques. We injected parkinsonian VGlut2-Cre mice with Cre-dependent channelrhodopsin (ChR2) and implanted 16-channel optrode arrays in the STN (Fig. 7A, Supplementary Fig. 6A). We then assessed STN single-unit firing during two optical stimulation paradigms in freely moving parkinsonian mice: “constant” and “pulsatile” (50 Hz) blue light stimulation (both 3 mW). As has been observed previously in anesthetized STN recordings in rats (Yu et al., 2020), STN neurons showed both excitatory and inhibitory responses to optical stimulation (Fig. 7B-E; Supplementary Fig. 6B-C). However, both stimulation paradigms caused similar decreases in overall firing rate (Fig. 7C,E; *P*=0.98 comparing relative firing rates during constant and 50 Hz stimulation, Wilcoxon sign-rank test). Of note, the response of STN neurons to optical stimulation differed between *in vivo*/freely moving and *ex vivo* preparations. During cell-attached patch-clamp recordings of STN in *ex vivo* slices (Supplementary Fig. 6D), pulsatile (50 Hz) blue light stimulation increased STN firing fairly consistently (Supplementary Fig. 6E-F; *P*=0.033), while constant illumination evoked more variable changes in spiking (Supplementary Fig. 6G-H; *P*=0.63). The effects of optical stimulation on STN firing in the *in vivo*, freely moving condition, however, are most relevant to the behavioral impact of STN DBS.

**Figure 7.**
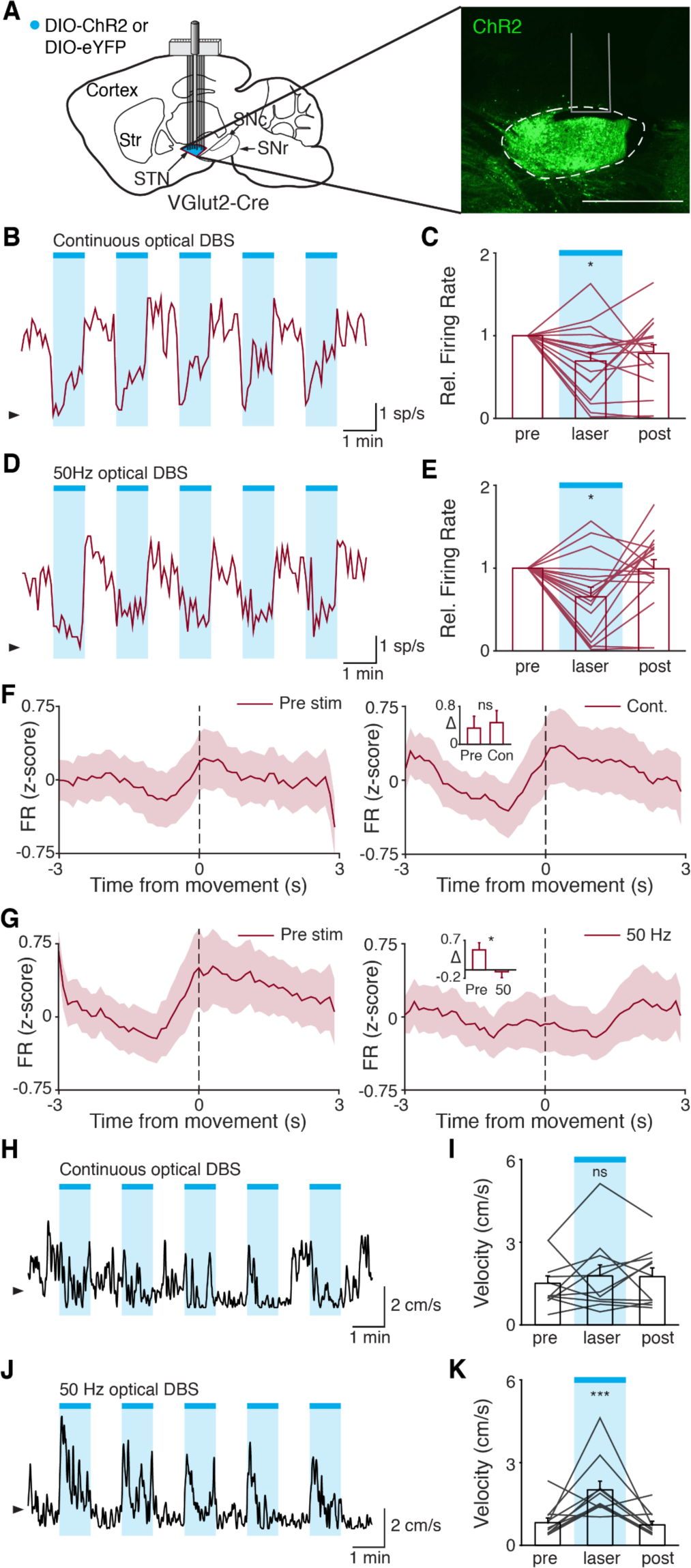
Disruption of STN dynamics is sufficient to provide therapeutic benefit. **(A)** Left: Sagittal schematic showing viral injection and 16-channel optrode implantation in the STN. Right: Postmortem sagittal section showing ChR2 expression in the STN (inset, scale=500 μm). **(B)** Representative STN single-unit firing in response to continuous optical stimulation, given in 1-minute epochs (blue bars). **(C)** Average z-scored firing rate of STN single units before, during, and after continuous optical stimulation (n=17 neurons, N=3 mice). **(D)** Representative STN single-unit firing in response to 50 Hz optical stimulation, given in 1-minute epochs (blue bars). **(E)** Average z-scored firing rate of STN single units before, during, and after 50 Hz optical stimulation (n=17 neurons, N=3 mice). **(F)** Average z-scored firing rate of STN single units aligned to movement starts before (left) and during (right) continuous optical stimulation. Inset: average change in firing rate around movement starts (see Methods for further details) before (pre) or during (con) constant optical stimulation (n=17 neurons, N=3 mice). **(G)** Average z-scored firing rate of STN single units aligned to movement starts before (left) and during (right) 50 Hz optical stimulation. Inset: average change in firing rate around movement starts before (pre) and during (50) 50 Hz optical stim (n=17 neurons, N=3 mice). **(H)** Representative single-session movement velocity in response to continuous optical stimulation, given in 1-minute epochs. **(I)** Average velocity before, during, and after continuous optical stimulation (N=11 mice). **(J)** Representative single-session movement velocity in response to 50 Hz optical stimulation, given in 1-minute epochs. **(K)** Average velocity before, during, and after 50 Hz optical stimulation (N=11 mice). Statistical significance was determined using a Wilcoxon rank-sum test (F-G) or a one-way repeated measures ANOVA with a Tukey HSD post hoc analysis applied to correct for multiple comparisons (C,E,I,K); *P<.05, \*\*\**P* < 0.001 (For ANOVA, only comparison between pre and laser shown, see Supplementary Table 1 for detailed statistics). Arrowhead in firing rate and velocity traces corresponds to 0 sp/s and 1 cm/s, respectively. Bar plots and average firing rate traces show mean ± SEM.

While both optical stimulation patterns produced decreases in the average firing rate of STN neurons, the two stimulation paradigms had different effects on movement-related STN dynamics. During stimulation-off epochs for both paradigms, STN firing rate increased around the time of movement starts (Fig. 7F,G, left), as we had observed previously (Fig. 1J). During constant optical stimulation, these movement-related STN dynamics did not change significantly (Fig. 7F, right; *P*=0.51). In contrast, pulsatile 50 Hz stimulation greatly attenuated these dynamics (Fig. 7G, right; *P*=0.039). Based on these electrophysiological recordings, continuous and 50 Hz optical stimulation produce similar changes in firing rate, but distinct effects on dynamics, allowing us to disentangle the effects of STN rate and pattern in producing therapeutic effects in parkinsonian mice.

To test whether these two stimulation paradigms would produce different behavioral effects, we studied a larger cohort of parkinsonian VGlut2-Cre mice, injected with Cre-dependent ChR2 or eYFP (control) and implanted with optical fibers in the STN (Supplementary Fig. 6A). In different test sessions, we stimulated with either constant or pulsatile blue light in 1-minute epochs. We predicted that continuous stimulation, which did not alter STN dynamics, would not produce benefits, while pulsatile stimulation, which greatly attenuated STN dynamics, would increase movement velocity. During constant illumination, neither STN-ChR2 nor STN-eYFP mice showed a significant increase in movement velocity or other metrics such as percent time moving or change in rotation bias (Fig. 7H-I; Supplementary Fig. 6I-K; *P*=0.30 for ChR2 movement velocity, *P*=0.66 for eYFP movement velocity). However, STN-ChR2 mice receiving 50 Hz stimulation did show a significant increase in movement velocity and percent time moving (Fig. 7J-K; Supplementary Fig. 6L-M; *P*=1.27x10^-8^ for movement velocity, *P*=5.56x10^-9^ for percent time moving), similar to mice receiving electrical STN DBS, and also developed an increased bias towards ipsilesional rotations (Supplementary Fig. 6L (inset); *P*=0.016). STN-eYFP mice did not show a significant change in movement velocity during 50 Hz stimulation (Supplementary Fig. 6N; *P*=0.81). These results demonstrate that although therapeutic electrical and optical STN DBS produce quite distinct changes on the rate of STN activity, they both disrupt movement-related STN dynamics. Moreover, they suggest that disrupting movement-related STN dynamics is sufficient to ameliorate parkinsonian motor deficits, making it a candidate mechanism for electrical STN DBS.

## Discussion

We combined a recently developed mouse model of electrical STN DBS for Parkinson’s disease with electrical artifact-free GCaMP fiber photometry to investigate three major theories surrounding the mechanism of electrical STN DBS: (1) STN DBS inhibits STN and SNr activity; (2) STN DBS acts through antidromic activation of hyperdirect M1 neurons; and (3) STN DBS acts through disrupting neuronal activity patterns within the STN. We observed that STN DBS excites rather than inhibits the STN and SNr, and that M1 activation does not seem to be necessary for therapeutic STN DBS, rendering the first two theories less likely. In support of the third theory, we observed that therapeutic electrical stimulation abolished patterned STN activity around movement onset and used optical manipulations to demonstrate that attenuating STN dynamics may be causal in the motor benefits of DBS.

In this study, we used calcium imaging with GCaMP6s to examine how neural activity changes during STN DBS. A recent study demonstrated technical feasibility of using *in vivo* calcium imaging with electrical STN DBS (Trevathan et al., 2020), but ours is the first to use this approach to link DBS-mediated physiological changes to behavior in freely moving parkinsonian animals. Moreover, our study is one of the first to use fiber photometry in deep basal ganglia nuclei, and to compare these signals with *in vivo* single-unit electrophysiology. This approach had several advantages, as well as limitations. The key advantage was the ability to obtain recordings free from electrical artifacts. Artifacts had been a major obstacle in prior electrophysiological studies, particularly in studying the effect of DBS on neurons in the target structure, such as the STN or GPi. A second advantage was the ability to use cell type- or projection-specific techniques. For example, the use of retrograde viruses allowed us to target the direct projections from primary motor cortex to STN (hyperdirect pathway), a population of significant interest in PD and DBS. Though STN and SNr are relatively homogeneous structures, with regard to major neurotransmitters (Smith and Parent, 1988; Walaas and Fonnum, 1979), future studies could use GCaMP and genetics to target either specific STN/SNr projections, or novel cell types within them (Kita and Kitai, 1987; Liu et al., 2020). A disadvantage of our approach over electrophysiology, however, is its temporal resolution. While traditional single-unit electrophysiology can detect individual action potentials, calcium imaging with GCaMP provides an integrated signal arising from multiple spikes (Chen et al., 2013; Sabatini, 2019). Fiber photometry further averages across a population of neurons, making it challenging to detect more rapid events. Given this temporal lag, it is difficult to establish whether DBS-associated changes in GCaMP signal arise directly from electrical stimulation, network activation, sensory feedback, or all of the above. Nonetheless, our activity measurements with photometry showed striking similarities to activity measurements using single-unit electrophysiology, over both short (movement-aligned activity) and long (responses to systemic levodopa administration) timescales. There thus appears to be correspondence between the two recording techniques. In the future, voltage indicators with high signal-to-noise that are compatible with deep imaging, as well as miniscope imaging of many single neurons simultaneously, may allow detection of single spikes and help increase the information obtained in optical recordings during DBS.

Our observation that therapeutic STN DBS increases activity at the level of the STN and SNr, rather than inhibiting it, is at odds with traditional rate-based models of basal ganglia function. In fact, previous STN DBS electrophysiological studies in parkinsonian primates have had conflicting results: some show increased basal ganglia output activity during STN DBS, while others observe local STN inhibition (Filali et al., 2004; Moran et al., 2011). Single-unit data in rodents is more limited, but analyses of pattern at both the single-unit and LFP level in one study suggest that the therapeutic benefit of STN DBS may be independent from overall changes in STN firing rate (Zhuang et al., 2018). Thus, it is possible that treatments that increase (e.g. DBS) or decrease STN firing (e.g. optical inhibition (Yoon et al., 2014) or levodopa) may be therapeutic. Some discrepancies in existing physiological data may arise from the imperfect process of removing STN DBS artifacts from electrophysiological recordings, especially in structures like STN and SNr that have high spontaneous firing rates. Others may relate to differences among animal models of PD.

Electrical STN DBS may increase the activity of STN neurons via several physiological mechanisms. As electrical stimulation has previously been suggested to preferentially recruit axons (McIntyre et al., 2004; McIntyre and Grill, 1999; Nowak and Bullier, 1998), DBS may drive antidromic spiking of STN neurons via stimulation of efferent STN axons, as well as drive STN spiking via activation of incoming excitatory axons. Depending on the proximity of the electrode to neural elements, it may drive STN activity through local activation of dendrites or cell bodies, as well. Increased SNr spiking may be driven by orthodromic activation of STN to SNr excitatory axons, or potentially by more complex network effects. The time resolution of fiber photometry and the time lag between STN and SNr activation (on average about 1 second) with this method cannot exclude polysynaptic effects. It is important to note that calcium imaging data are correlative, and the behavior of mice during therapeutic STN DBS changes markedly. The change in movement may itself drive changes in STN and SNr activity, for example through sensory feedback to the basal ganglia. The recorded signals in the STN and SNr may therefore be both a cause and/or an effect of increased movement.

Though we observed modulation of hyperdirect M1 neurons during STN DBS, this modulation did not correlate well with therapeutic effects. While this is suggestive that the hyperdirect pathway is less important for the therapeutic effect of STN DBS, differences in the physiologic properties of M1 projection neurons, as compared to STN and SNr neurons, might have contributed to the late-developing changes in M1. However, removal of M1 did not abolish the therapeutic benefit of STN DBS, leading us to conclude that hyperdirect M1 activation may not play a central role. The discrepancy between our conclusions and those of previous studies has a number of potential explanations. While past mouse studies have used optogenetic stimulation as a proxy for electrical STN DBS (Gradinaru et al., 2009; Sanders and Jaeger, 2016), we used electrical stimulation in an effort to more closely model what is observed in PD patients (our eventual use of optogenetic stimulation was directly informed by our observations during electrical stimulation). The former approach identifies manipulations that are sufficient to relieve parkinsonian motor symptoms, while the latter identifies changes that correlate with a specific therapy. Thus, while optogenetic stimulation may reveal that changing neural activity in a variety of ways can relieve parkinsonism in mice, it is difficult to extrapolate which of these changes actually occur during electrical STN DBS. In other words, two therapies that have similar behavioral effects may not have the same mechanism of action. In addition, while hyperdirect pathway neurons did not seem to be crucial for the benefits of STN DBS in mice, using locomotor velocity as a primary outcome measure, they may play an important role in other species, or in the benefits of DBS in other motor domains.

In the absence of therapeutic manipulations, we found that STN neurons in parkinsonian mice show an increase in activity around movement initiation. Similar observations have been made in the STN of healthy NHPs and cats (Cheruel et al., 1996; Wichmann et al., 1994). Less is known about what drives this increase, though candidates include excitatory input from hyperdirect M1 neurons (Polyakova et al., 2020), inhibitory input from the globus pallidus pars externa (GPe) (Chu et al., 2015; Kovaleski et al., 2020), and/or increased synchronization among STN neurons. As we found that disruption of STN movement-related dynamics may play a role in alleviating parkinsonian symptoms, identifying the physiological sources of this signal will likely be an important question in future research.

Our observation that movement-related STN activity patterns are altered during STN DBS may relate to previous work showing changes in firing patterns or network synchrony during STN DBS. It has previously been postulated that rate-independent aspects of neural activity, such as within-neuron firing pattern or between-neuron synchronization, may drive PD symptoms and represent key markers of therapeutic interventions (Hammond et al., 2007; Little and Brown, 2014; Wichmann, 2019). Many other groups have observed increased oscillations throughout the basal ganglia in parkinsonian animal models and in humans, which may resolve with therapeutic treatment (de Hemptinne et al., 2015; Halje et al., 2012; Moran et al., 2012; Shimamoto et al., 2013). In fact, one group studying healthy NHPs has even observed positive modulation during movement in pallidal cells, similar to what we observed during movement starts in STN neurons, and which was similarly interrupted in a subset of pallidal neurons during STN DBS (Zimnik et al., 2015). The difficulty, though, has been in establishing a causal link between changes in these patterns during DBS and improvement in behavior. Our ability to not only observe, but also to trigger these changes through optogenetic manipulation helps fill this critical gap. Additional causal links might be investigated further in the future using a combination of optical and electrical methods, building on the approach introduced here. Though it facilitated our use of cell type-specific methods and imaging tools, a key potential caveat of our study is the mouse model of PD. First, the 6-OHDA model causes focal, rapid, and in this case nearly complete loss of dopaminergic neurons and their projections. In contrast, PD causes neurodegenerative changes in multiple brain areas over many years. Though many key physiological features of PD are similar in toxin-based models of parkinsonism (Bové and Perier, 2012; Campos et al., 2013), others may be distinct based on the pattern and tempo of neurodegeneration. In addition, electrical stimulation may have different impacts in the small mouse brain as compared to the human brain. Though electrical stimulation is unlikely to respect the borders of brain nuclei in either species, the small size of the target region (STN) in the mouse brain increases the likelihood that fibers in adjacent areas are recruited by DBS. Comfortingly, our observation that the therapeutic effects of electrical STN DBS in the mouse fall off rapidly below the average current amplitude (200 uA) and vary according to the STN region targeted (Schor and Nelson, 2019) match well with human data and suggest some specificity in the volume of tissue activated (Greenhouse et al., 2011; Tommasi et al., 2008).

For practical reasons, in order to test the physiological and behavioral effects of a wide variety of stimulation parameters (both subtherapeutic and therapeutic), we delivered short (1-minute) epochs of stimulation. These short epochs may not capture the longer term changes in neural activity expected in PD patients, where continuous high frequency stimulation is the current clinical standard. Reducing this concern, many DBS benefits indeed evolve rapidly in PD patients, as in the mouse model (Hristova et al., 2000; Temperli et al., 2003), and our previous work demonstrates consistent behavioral improvement in the model even across longer timescales (Schor and Nelson, 2019).

Excitingly, our observation that non-canonical changes in STN activity confer therapeutic benefit in a mouse model of PD suggests a wider therapeutic space for the treatment of PD. Many therapeutic approaches to PD have been predicated on the idea that inhibition of hyperactive basal ganglia nuclei is required for therapeutic benefit, but our findings, as well as recent work using close-loop DBS (Bouthour et al., 2019; Johnson et al., 2016; Rosin et al., 2011) indicate non-rate-based alterations in activity can improve movement. In addition, our work linking neural activity to behavior in STN DBS for PD may inform the application of DBS to other neuropsychiatric disorders. To rationally apply DBS to other conditions, such as addiction or Tourette’s syndrome, it is critical to know how electrical stimulation might impact the underlying neural circuitry of disease. We hope that our work may serve as a blueprint for future inquiries into the therapeutic potential of DBS.

## Materials and Methods

### Animals

3-6 month-old wild-type and transgenic C57Bl/6 mice of either sex were used in this study. To allow optical recording and manipulation of glutamatergic STN neurons, homozygous VGlut2-Cre mice (Stock No. 028863, Jackson Labs) were bred to wild-type C57BL**/**6 mice (Jackson Labs) to yield hemizygous VGlut2-Cre mice. To allow optical recording of GABAergic SNr neurons, homozygous VGAT-Cre mice (Jackson Labs) were bred to wild-type C57BL**/**6 mice (Jackson Labs) to yield hemizygous VGAT-Cre mice. Animals were housed 1-5 per cage on a 12-hour light/dark cycle with *ad libitum* access to rodent chow and water. All behavioral manipulations were performed during the light phase. We complied with local and national ethical regulations regarding the use of mice in research. All experimental protocols were approved by the UC San Francisco Institutional Animal Care and Use Committee.

### Electrical DBS Devices

We constructed electrical DBS devices consisting of 3 twisted pairs of stainless steel wire (76.2 micron diameter, coated, AM Systems), cut at an angle to span approximately 300 microns in DV. These were pressure-fit into female Millmax connectors. Each electrode pair was tested for shorts prior to electrode implantation. For additional details see Schor and Nelson, 2019.

### Surgical Procedures

Stereotaxic surgery was performed between 3 and 6 months of age. Anesthesia was induced with intraperitoneal (IP) injection (0.1 mL) of ketamine (40 mg/kg) and xylazine (10 mg/kg) and maintained with inhaled isoflurane (0.5%-1%). To model Parkinson’s disease in mice, the neurotoxin 6-hydroxydopamine (6-OHDA, 1 μL, 5 mg/mL) was injected unilaterally in the left medial forebrain bundle (MFB, -1.0 AP, -1.0 ML, 4.9 DV from Bregma). Desipramine (0.2 mL, 2.5 mg/mL) was injected intraperitoneally (IP) approximately 30 min prior to 6-OHDA injections to reduce uptake by other monoaminergic neurons in the MFB. Additional surgeries were performed at least two weeks following 6-OHDA injection.

For experiments involving combined electrical STN DBS and optical imaging, a 3-lead bipolar stimulating electrode array was implanted in the ipsilesional STN (-1.8 AP, -1.65 ML, 4.5 DV) (Schor and Nelson, 2019). During the same surgery, VGlut2-Cre mice were injected with Cre-dependent AAV1-Syn-Flex-GCaMP6s-WPRE-SV40 (UPenn, 100 nL injected diluted 1:8 in normal saline, undiluted titer 3.06 x 10^13^/mL) in the STN (-1.8 AP, - 1.65 ML, 4.5 DV) and implanted with a photometry fiber-optic ferrule (0.4 mm, Doric Lenses) above the STN (4.3 DV). VGAT-Cre mice were injected with the same Cre-dependent GCaMP6s vector (300-500 nL injected diluted 1:8 in normal saline) in the SNr (-3.2 AP, -1.6 ML, 4.5 DV) and implanted with a fiber-optic ferrule above the SNr (4.3 DV). Wild-type mice were injected with a retrograde virus encoding Cre recombinase [either CAV-Cre (Montpellier, 100 nL injected undiluted, undiluted titer 1.0 x 10^13^/mL) or AAV2retro-Cre-mCherry (Addgene/UPenn Vector Core, 100 nL injected undiluted, undiluted titer 7.8 x 10^13^/mL)] in the STN (-1.8 AP, -1.65 ML, 4.5 DV) and Cre-dependent GCaMP6s (500 nL injected diluted 1:8 in normal saline) in the primary motor cortex (M1, +2 AP, -1.56 ML, 1 DV) and implanted with a fiber-optic ferrule above M1 (0.8 DV).

For *in vivo* electrophysiological experiments, mice were implanted with a 16-channel 7 mm fixed electrode array (Innovative Neurophysiology) in either the STN or the SNr, using the same coordinates as used for GCaMP6s injections above. For combined *in vivo* electrophysiology/optical stimulation experiments, VGlut2-Cre littermates were injected with Cre-dependent AAV5-DIO-ChR2-eYFP (UPenn, injected diluted 1:2 in normal saline, 100 nL, undiluted titer 1.02 x 10^13^/mL) or AAV5-DIO-eYFP (UNC, injected undiluted, 100 nL, titer 4.4 x 10^12^/mL) in a randomized fashion, and implanted with a 16-channel 7 mm fixed electrode array (Innovative Neurophysiology) with a fiber-optic ferrule (0.2 mm, Thor Labs) epoxied ∼0.2mm above the electrode tips in the STN. For optical stimulation experiments without recording, a fiber-optic ferrule was implanted just above the STN (4.3 DV). A minimum of 3 weeks were allowed for viral expression before behavioral testing.

For experiments involving M1 lesioning, a large rectangular craniectomy was performed to expose brain tissue containing M1 (vertices of rectangle at [-0.1 AP, -2.1 ML]; [2.6 AP, -2.1 ML]; [2.6 AP, -0.9 ML]; and [-0.1 AP, -0.9 ML]). A micro knife (FST) was then used to carefully remove a 1 mm thick rectangle of brain tissue that was the height and width of the craniectomy under a dissecting microscope. A hemostatic sponge (Ethicon) was used to staunch any bleeding before covering the lesion and adjacent bone with silicone sealant (Kwik-Cast). A 3-lead bipolar stimulating electrode array was then implanted in the ipsilesional STN as previously described. A minimum of 10 days of recovery was allowed before subsequent behavioral testing.

### Behavior

All behavior was conducted in the open field (clear acrylic cylinders, 25 cm diameter) following 1 day of habituation (20 minutes). Mice were monitored via two cameras, one directly above and one in front of the chamber. Video-tracking software (Noldus Ethovision) or custom-written code (Matlab) was used to quantify locomotor activity, including movement velocity, ipsilateral rotations, and contralateral rotations. Dyskinesia was scored manually by an unblinded rater using a modified version of the abnormal involuntary movement (AIM) scoring method (Cenci and Lundblad, 2007). Dyskinesia was quantified in one-minute increments either every minute (for STN DBS experiments) or every 5 minutes (for levodopa experiments), with axial, limb, and orofacial body segments rated on a scale of 0-3 each. A score of 0 indicates no abnormal movement, while a score of 3 indicates continuous dyskinesia for the one-minute epoch. The scores for each body segment are then summed, with a maximum score of 9 per epoch.

### Pharmacology

6-OHDA (Sigma Aldrich) was prepared at 5 mg/mL in normal saline. Levodopa was prepared (0.5 mg/mL Sigma Aldrich) with benserazide (0.25 mg/mL, Sigma Aldrich) in normal saline and always administered at 5 mg/kg.

### Electrical Stimulation

An isolated constant current bipolar stimulator (WPI) was used to deliver electrical stimuli. The timing of stimuli was controlled by TTL input from an Arduino. Electrical stimulation experiments consisted of five 1 min stimulation periods, each preceded and followed by 1 min of no stimulation, for a total of 11 min. Both the construction of STN DBS electrodes and the determination of optimal stimulation electrode pair were as detailed previously (Schor and Nelson, 2019).

### Fiber Photometry

Fiber photometry signals were acquired through implanted 400 *μ*m optical fibers, using an LED driver system (Doric). Following signal modulation, 405 nm (control signal, from GCaMP autofluorescence) and 465 nm signals were demodulated via a lock-in amplifier (RZ5P, TDT), visualized, and recorded (Synapse, TDT). Offline, the 405 nm signal was fit to the 465 nm signal using a first-degree polynomial fit (Matlab) to extract the non-calcium dependent signal (due to autofluorescence, fiber bending, etc). The fitted 405 nm signal was then subtracted from the 465 nm signal to generate a motion-corrected signal. Animals in which the fitted 405 nm signal did not differ from the 465 nm signal were excluded from further analysis. To remove the gradual, slow bleaching observed in the ∼3 hour saline and levodopa recordings, we additionally fit a double exponential to the 405 nm signal, linearly fit it to the the motion-corrected signal, and then subtracted it.

Every processed fiber photometry signal was normalized (z-scored) by subtracting the mean and dividing by the standard deviation of the closest preceding “pre” period. For electrical stimulation experiments, the 30 seconds preceding each stimulation period was used to normalize the subsequent 1-min stim and 1-min post period. For levodopa and saline experiments, the 20 minutes prior to injection was used to normalize the subsequent 2.5 hours of signal.

### *In vivo* Electrophysiology

Single-unit activity from microwires was recorded using a commutated (Doric) multiplexed 96-channel recording system (CerePlex Direct, Blackrock Microsystems). Spike waveforms were filtered at 154–8800 Hz and digitized at 30 kHz. The experimenter manually set a threshold for storage of electrical events. Spike sorting and single units (SUs) were identified offline by manual sorting into clusters (Offline Sorter, Plexon). Waveform features used for separating units were typically a combination of valley amplitude, the first three principal components (PCs), and/or nonlinear energy. Clusters were classified as SUs if they fulfilled the following criteria: (1) <1% of spikes occurred within the refractory period and (2) the cluster was statistically different (p<0.05, MANOVA using the aforementioned features) from the multi- and other single-unit clusters on the same wire.

### *Ex vivo* Slice Electrophysiology and Imaging

To prepare *ex vivo* slices for whole-cell recordings and GCaMP imaging, mice were deeply anesthetized with IP ketamine-xylazine, transcardially perfused with ice-cold glycerol-based slicing solution, decapitated, and the brain was removed. Glycerol-based slicing solution contained (in mM): 250 glycerol, 2.5 KCl, 1.2 NaH_2_PO_4_, 10 HEPES, 21 NaHCO_3_, 5 glucose, 2 MgCl_2_, 2 CaCl_2_. The brain was mounted on a submerged chuck, and sequential 275 mm coronal or sagittal slices were cut on a vibrating microtome (Leica), transferred to a chamber of warm (34°C) carbogenated ACSF containing (in mM) 125 NaCl, 26 NaHCO_3_, 2.5 KCl, 1 MgCl_2_, 2 CaCl_2_, 1.25 NaH2PO4, 12.5 glucose for 30-60 min, then stored in carbogenated ACSF at room temperature. Each slice was then submerged in a chamber superfused with carbogenated ACSF at 31°C-33°C for recordings. STN neurons were targeted using differential interference contrast (DIC) optics in VGlut2-Cre mice on an Olympus BX 51 WIF microscope.

For opsin validation experiments, neurons were patched in the cell-attached configuration using borosilicate glass electrodes (3-5 MOhms) filled with ACSF. Picrotoxin was added to all external solutions for opsin validation. For combined electrophysiology-imaging experiments with GCaMP6s, neurons were patched in the whole-cell current-clamp configuration using borosilicate glass electrodes (3-5 MOhms) filled with potassium methanesulfonate-based internal solution containing (in mM): 130 KMeSO_3_, 10 NaCl, 2 MgCl_2_, 0.16 CaCl_2_, 0.5 EGTA, 10 HEPES, 2 MgATP, 0.3 NaGTP, pH 7.3. All recordings were made using a MultiClamp 700B amplifier (Molecular Devices) and digitized with an ITC-18 A/D board (HEKA). Data were acquired using Igor Pro 6.0 software (Wavemetrics) and custom acquisition routines (mafPC, courtesy of M. A. Xu-Friedman). Recordings were filtered at 5 kHz and digitized at 10 kHz.

To validate ChR2 function in slice, light pulses were delivered to the slice by a TTL-controlled LED (Olympus), passed through a GFP (473 nm) filter (Chroma) and the 40X immersion objective. LED intensity was adjusted to yield an output of 3 mW at the slice. Light was delivered in 1-minute epochs, at 50 Hz, 3 ms pulse width or continuously. Stimulation lasted for 1 min and was preceded and followed by 30 seconds of recording without stimulation.

For simultaneous electrophysiology and GCaMP6s imaging, current-clamped neurons were stimulated (0.5-1 nA) to elicit action potentials. Stimulation occurred at 10, 20, 40, 50 or 60, and 100 or 120 Hz (100 μs pulse-width); or was delivered as a long single square wave of constant current for 1 min, preceded and followed by 30 seconds without stimulation. During the duration of each 2 min trial GCaMP fluorescence was either acquired through 1-photon or 2-photon microscopy. 1-photon experiments used a 473 nm light (TTL-controlled LED, Olympus, paired with GFP filter, Chroma) delivered to the slice at <1 mW, with GCaMP6s fluorescence captured using an imaging camera attached to the microscope (QI Retiga Electro). For 2-photon microscopy, a 2-photon source (Coherent Ultra II) was tuned to 810 nm to identify GCaMP expressing neurons and tuned to 940 nm for calcium imaging. Epi- and transfluorescence signals were captured through a 40×, 0.8 NA objective paired with a 1.4 NA oil immersion condenser (Olympus) to photomultiplier tubes (H10770PA-40 PMTs, Hamamatsu). Data were collected in line scan mode (2–2.4 ms/line, including mirror flyback).

All *ex vivo* electrical recordings were passed through a 1 Hz high-pass filter to remove slow electrical drift and spikes were extracted using the findpeaks function in Matlab. All *ex vivo* optical recordings were first collapsed into a one-dimensional fluorescence time series by averaging the fluorescence of pixels within a defined region-of-interest. In one-photon recordings, this signal was further processed by fitting a double exponential and subtracting it to remove effects of signal bleaching.

### Optogenetic Manipulations

Prior to optical stimulation experiments, animals were habituated to tethering with custom lightweight patch cables (Precision Fiber Products and ThorLabs) coupled to an optical commutator (Doric Lenses) in the open field for 30 min per day, over 1-2 days. Optical stimulation sessions consisted of five 1 min stimulation periods, each preceded and followed by 1 min of no stimulation, for a total of 11 min. TTL-controlled (Master8, A.M.P.I.) blue laser light (488 nm, 3 mW, Shanghai Laser and Optics Century) was delivered in pulse trains (3 ms, 50Hz) or continuously. Behavior was rated by an observer blinded to the injected construct (ChR2-eYFP or eYFP).

### Histology and Microscopy

Mice were terminally anesthetized with IP ketamine (200 mg/kg) and xylazine (40 mg/kg). For mice with an implanted STN DBS device or multielectrode array, the site of stimulation or recording was marked with a solid state, direct current Lesion Maker (Ugo Basile). Mice were then transcardially perfused with 4% paraformaldyde (PFA), the brain was dissected from the skull and fixed overnight in 4% PFA, and then was placed in 30% sucrose at 4°C for 2-3 days. Brains were then cut into 50 μm sagittal sections on a freezing microtome (Leica). To confirm dopamine depletion, tissue was immunostained for tyrosine hydroxylase (TH). Stitched multi-channel fluorescence images were taken on a Nikon 6D conventional widefield microscope at 4-10X, using custom software (UCSF Nikon Imaging Center) to confirm virus expression, fiber placement, and STN DBS placement.

### Group Allocation and Blinding

The order in which each mouse received electrical stimulation during optical recording experiments was randomized daily, as was the type of stimulation administered. Mouse order and stimulation type was also randomized during optogenetic experiments. For optical manipulations, VGluT-Cre positive littermates were randomized to eYFP or ChR2-eYFP injection. The experimenter was blinded to experimental group (eYFP vs ChR2-eYFP) during behavioral experiments.

### Inclusion/Exclusion Criteria

If the number of ipsilesional TH-positive SNc neurons were >5% of the contralateral (unlesioned) side, the animal was excluded from all analyses. Animals that did not show strong virus expression, proper optical fiber placement (within target structure or <0.2 mm above), and/or proper STN DBS device placement (within STN) were excluded from further analysis. If DBS leads developed a short, further DBS experiments were terminated, but levodopa experiments were continued.

A short developed in one mouse in the STN DBS/SNr imaging cohort, such that only levodopa experiments could be performed. We failed to deliver 1 of the 30 DBS parameter combinations in one mouse from the STN DBS/M1 hyperdirect imaging cohort, so no 100 Hz DBS data were included for this mouse. Across all imaging cohorts, 4 mice were excluded due to insufficient GCaMP expression and/or improper targeting of the optical fiber, resulting in no detectable GCaMP signal on fiber photometry (4 of 30 imaging mice). Across optical stimulation cohorts, 2 mice were excluded due to insufficient eYFP or ChR2-eYFP expression (2 of 22 optical stimulation mice). One mouse in the STN in vivo electrophysiology cohort (1 of 4) and one mouse from the SNr in vivo electrophysiology cohort (1 of 4) were excluded due to a lack of clearly isolated single units.

### Sample Size Determination, Quantification, Statistical Analysis, and Replication

For *in vivo* imaging experiments, no similar studies had been performed by which to estimate effect size. We performed small pilot experiments in each recorded brain region to determine the mean and standard deviation of GCaMP signals in these regions. Effect sizes were estimated from these pilots, or from similarly designed and published electrophysiological studies. The sample size was calculated using 0.90 power to detect a significant difference, two-sided nonparametric comparisons, and alpha of 0.05. Sample sizes for *ex vivo* experiments, *in vivo* electrophysiology, and *in vivo* optical stimulation were calculated using a similar approach, but based on variance and effect sizes from previous experiments conducted in the lab using similar methods as well as published studies from other laboratories.

All data are expressed as mean ± standard error of the mean (SEM). For all bar graphs for electrical stimulation, optogenetic, and slice experiments, the “stim” or ”laser” bar was calculated by averaging all one-minute stimulation periods for each trial. The “pre” and “post” bars were calculated by averaging the 30 seconds before and 30 seconds after each stimulation period, respectively. For all bar graphs involving levodopa or saline, the “LD” or “saline” bar was calculated by averaging the ten minutes between 30-40 min post injection for each trial. The “pre” and “post” bars were calculated by averaging the time period between 15 min and 5 min before injection and between 125 min and 135 min post injection, respectively, for each trial. Correlation between photometry signal and velocity (Supplementary Fig. 5E) was calculated using *fitlm* in Matlab. Rise time of velocity and calcium signals was calculated as the time it took from the onset of stimulation for the signal to first reach the mean value for that stimulation epoch. Movement starts were defined as events when the mouse’s movement velocity changed from less than 0.5 cm/s (maintained for at least 1 s) to more than 2 cm/s. Change in photometry signal or firing rate around movement starts (insets in Fig. 6D and Fig. 7F-G) were calculated by subtracting the average fiber signal or firing rate during 1 sec preceding movement start from the average fiber signal or firing rate during 1 sec following movement start.

All data on which a repeated measures one-way ANOVA (rmANOVA) was performed were tested for normality using a Kolmogorov-Smirnov (KS) test. Nonparametric tests (Friedman test, Wilcoxon sign-rank, Wilcoxon rank-sum) were used in all other cases (see Supplementary Table 1 for full details). For rmANOVAs and Friedman tests, a Tukey HSD post hoc analysis was applied to correct for multiple comparisons. Data was considered statistically significant for p<0.05.

Multiple cohorts were used in each experiment, and findings were reliably reproduced among all subjects. *In vivo* optical recordings for each brain region were conducted using at least 3 cohorts of animals. Optogenetic experiments were conducted with 3 cohorts of animals. Electrophysiology experiments were conducted with at least 2 cohorts of animals. M1 lesion experiments were conducted with 2 cohorts of animals.

## Acknowledgements

The authors would like to acknowledge M. McGregor, P. Starr, J. Ostrem, G. Bouvier, M. Scanziani, and members of the Nelson and Bender Labs for providing advice and feedback on the manuscript. This work was supported by the NINDS (K08 NS081001, ABN; R01, ABN; F31 NS110329, JSS) and the UCSF Discovery Fellows Program (JSS). ABN is the Richard and Shirley Cahill Endowed Chair in Parkinson’s Disease Research.

## Author Contributions

JSS, ABN, and KJB designed the experiments. JSS, IGM, PWES, ABN, and KJB performed experiments. JSS, RJB, and JAS performed histology. JSS and ABN wrote the manuscript with contributions from all authors.

