## Supplemental Figures and Tables for "Therapeutic Deep Brain Stimulation Disrupts Subthalamic Nucleus Activity Dynamics in Parkinsonian Mice"

#### Supplementary Figures

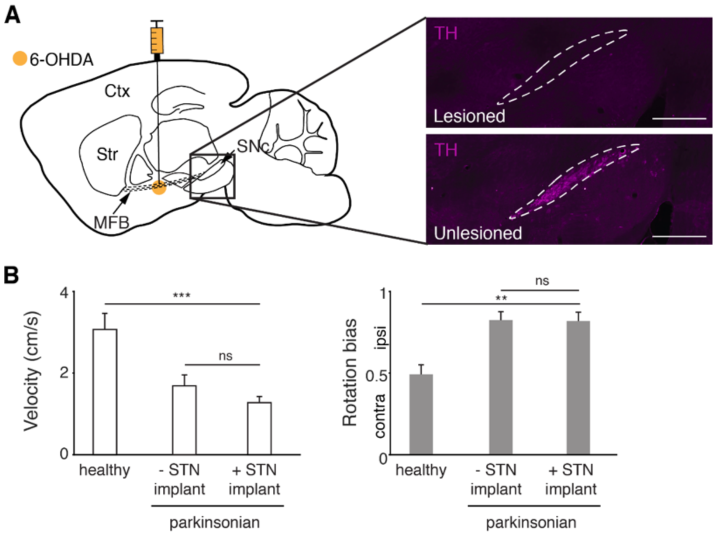


**Supplementary Figure 1. Related to Figure 1. Hemiparkinsonian mice show decreased movement velocity and ipsilesional rotation bias**. **(A)** Left: Sagittal schematic showing unilateral injection of 6-OHDA to deplete ipsilateral dopamine neurons. Right: Postmortem sagittal section showing depletion of TH (purple) in lesioned hemisphere (dotted lines indicate borders of substantia nigra, pars compacta; scale bar=500 µm). **(B)** Comparison of average velocity (left) and rotation bias (right) in healthy mice (N=9 mice), hemiparkinsonian mice without STN implants (N=10 mice), and hemiparkinsonian mice with STN implants (N=9 mice). Statistical significance was determined using a Wilcoxon rank-sum test; **P<.01, ****P*<0.001 (see Supplementary Table 1 for detailed statistics). Bar plots show mean ± SEM.

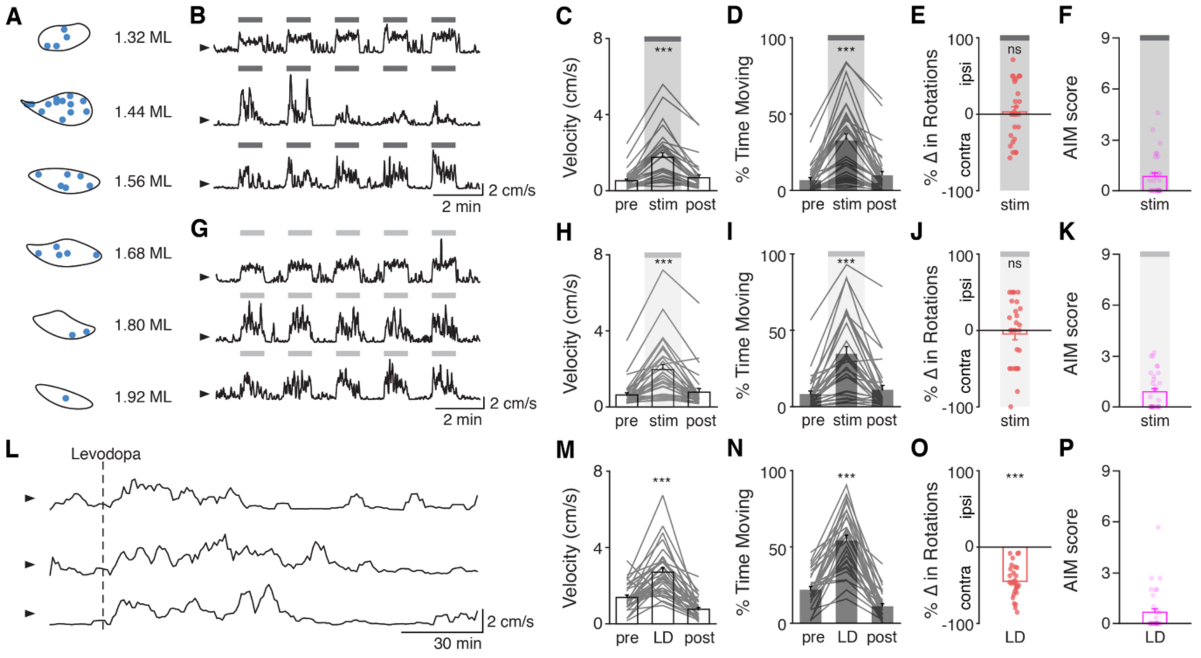


**Supplementary Figure 2. Related to Figures 2-4. STN DBS and levodopa produce similar behaviors in parkinsonian mice**. **(A)** Targeting of STN DBS electrodes to the STN in 32 mice. **(B)** Representative single-session velocities in response to 60 Hz STN DBS in 3 mice. **(C)** binned average velocity, **(D)** percent time moving, **(E)** change in rotational bias, and **(F)** dyskinesia in response to 60 Hz STN DBS (N=32 mice). **(G)** Representative single-session velocities in response to 100 Hz STN DBS in 3 mice. **(H)** binned average velocity, **(I)** percent time moving, **(J)** change in rotational bias, and **(K)** dyskinesia in response to 100 Hz STN DBS (N=31 mice). **(L)** Representative single-session velocities before and after levodopa injection (dotted line) in 3 mice. **(M)** binned average velocity, **(N)** percent time moving, **(O)** change in rotational bias, and **(P)** dyskinesia in response to levodopa injection (N=30 mice). Dyskinesia was quantified with as the abnormal involuntary movement (AIM) score. Statistical significance was determined using a Wilcoxon sign-rank test (E,I,O); a one-way repeated measures

ANOVA with a Tukey HSD post hoc analysis applied to correct for multiple comparisons (M-N); or a Friedman test with a Tukey HSD post hoc analysis applied to correct for multiple comparisons (C,D,H,I); ***$P<0.001$ (For ANOVA/Friedman, only comparison between pre and stim/LD shown, see Supplementary Table 1 for detailed statistics). Arrowhead in velocity traces corresponds to 1 cm/s. Bar plots show mean ± SEM.

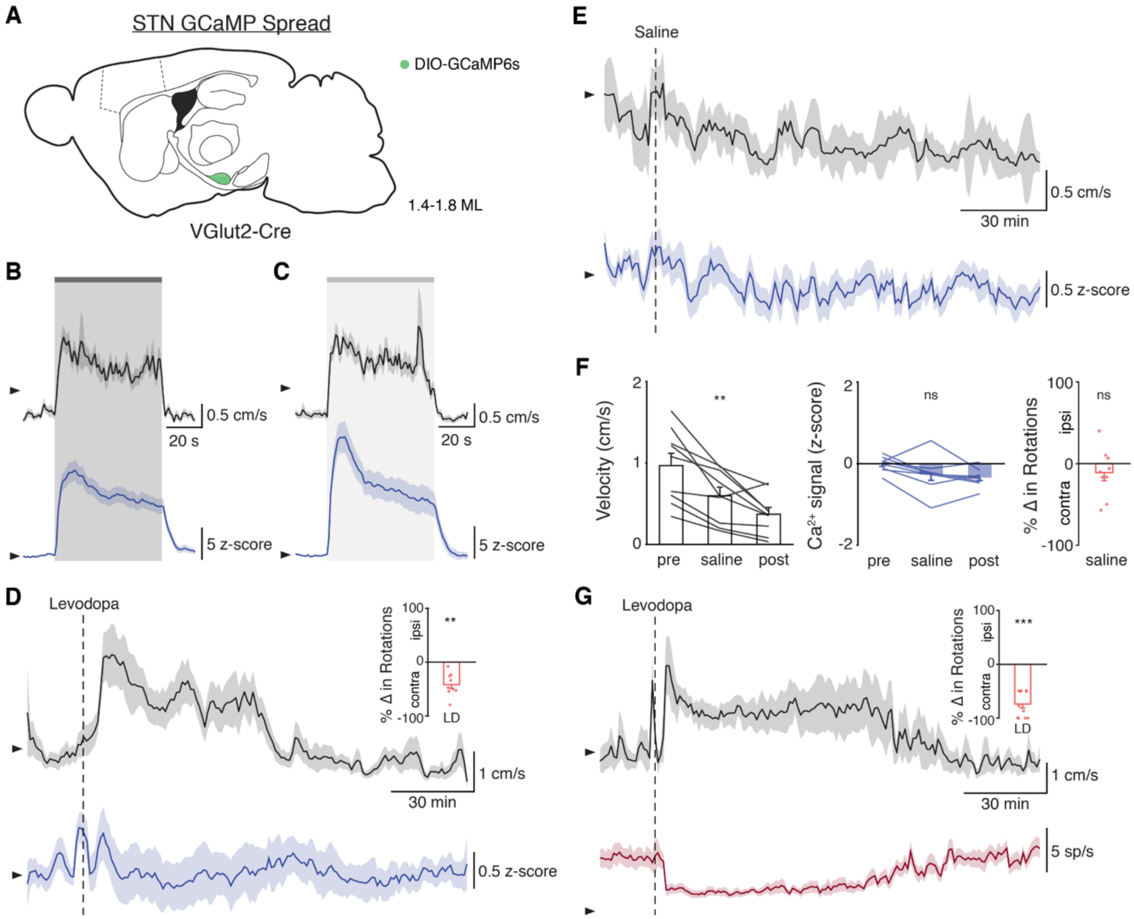


**Supplementary 3. Related to Figure 2. STN stimulation increases STN activity *in vivo*. (A)** Sagittal schematic showing estimated extent of viral GCaMP6s spread in the STN of VGlut2-Cre mice (N=9 mice). **(B-C)** Average movement velocity over time (black) and STN GCaMP signal (blue) following 60 Hz STN DBS (a, N=9 mice) or 100 Hz STN DBS (b, N=9 mice). **(D)** Average movement velocity over time (black) and STN GCaMP signal (blue) following administration of levodopa (N=9 mice) and average change in rotation bias during levodopa (inset). **(E)** Average movement velocity over time (black) and STN GCaMP signal (blue) following administration of saline (N=9 mice). **(F)** Average velocity (left), STN GCaMP signal (middle), and change in rotation bias before, during, and after saline injection (N=9 mice). **(G)** Average velocity over time (black) and STN single-unit firing rate (red) following administration of levodopa (n=11 cells, N=3 mice) and average change in rotation bias during levodopa (inset). Statistical significance was determined using a Wilcoxon sign-rank test (D,F (right),G) or a one-way repeated measures ANOVA with a Tukey HSD post hoc analysis applied to correct for multiple comparisons (F (left, middle)); **$P<0.01$, ***$P<0.001$ (For ANOVA, only comparison between pre and saline shown, see Supplementary Table 1 for detailed statistics). Arrowhead in velocity, GCaMP, and single-unit electrophysiology traces corresponds to 1 cm/s, 0 z-score, and 0 sp/s, respectively. Velocity traces, GCaMP traces, single-unit electrophysiology traces, and bar plots show mean ± SEM.

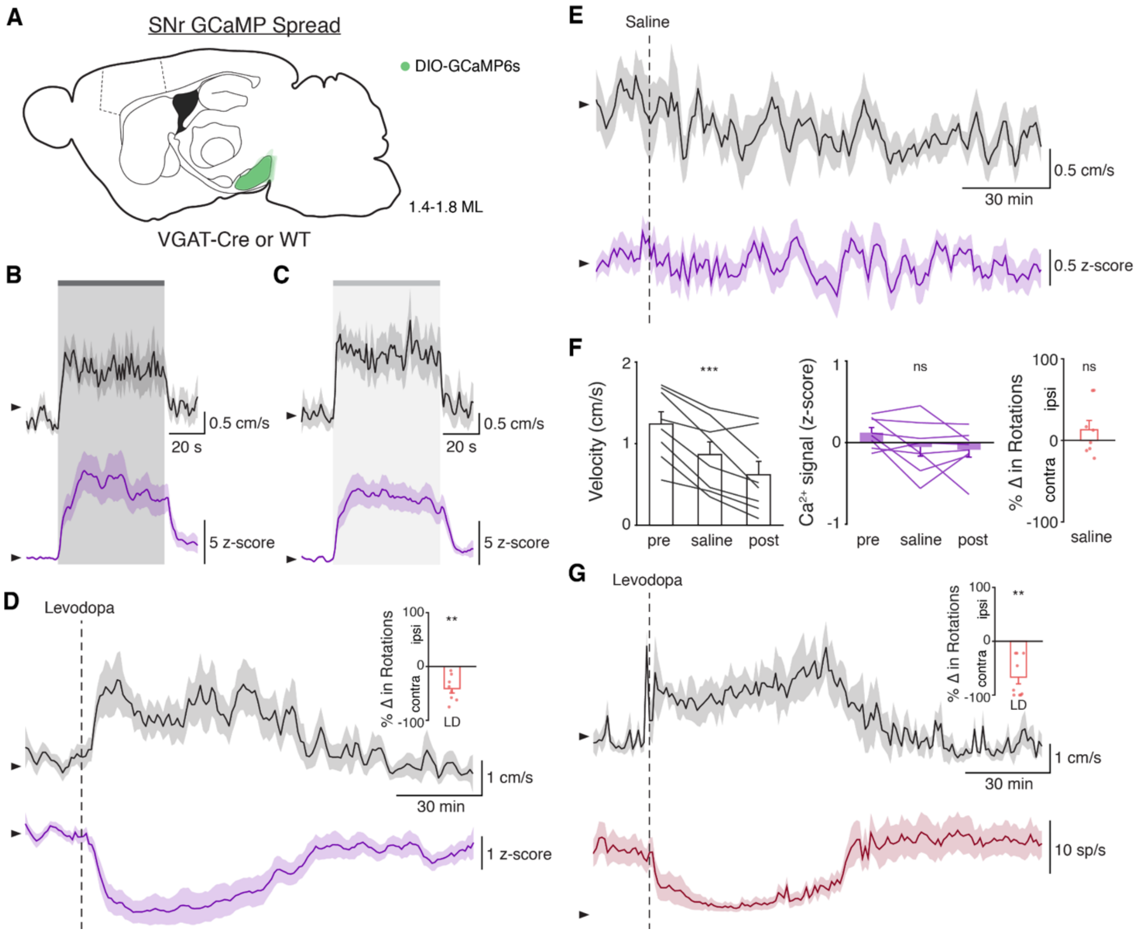


**Supplementary Figure 4. Related to Figure 3. STN DBS evokes a rapid increase in SNr activity. (A)** Sagittal schematic showing estimated extent of viral GCaMP6s spread in the SNr of VGAT-Cre or WT mice (N=8 mice). **(B-C)** Average velocity over time (black) and SNr GCaMP signal (purple) following 60 Hz STN DBS (a, N=7 mice) or 100 Hz STN DBS (b, N=7 mice). **(D)** Average velocity over time (black) and SNr GCaMP signal (purple) following administration of levodopa (N=8 mice) and average change in rotation bias during levodopa (inset). **(E)** Average velocity over time (black) and SNr GCaMP signal (purple) following administration of saline (N=8 mice). **(F)** Average velocity (left), SNr GCaMP signal (middle), and change in rotation bias before, during, and after saline injection (N=8 mice). **(G)** Average velocity over time (black) and SNr single-unit firing rate (red) following administration of levodopa (n=9 cells, N=3 mice) and average change in rotation bias during levodopa (inset). Statistical significance was determined using a Wilcoxon sign-rank test (D,F (right),G) or a one-way repeated measures ANOVA with a Tukey HSD post hoc analysis applied to correct for multiple comparisons (F (left, middle)); $^{**}P<0.01$, $^{***}P<0.001$ (For ANOVA, only comparison between pre and saline shown, see Supplementary Table 1 for detailed statistics). Arrowhead in velocity, GCaMP, and single-unit electrophysiology traces corresponds to 1 cm/s, 0 z-score, and 0 sp/s, respectively. Velocity traces, GCaMP traces, single-unit electrophysiology traces, and bar plots show mean ± SEM.

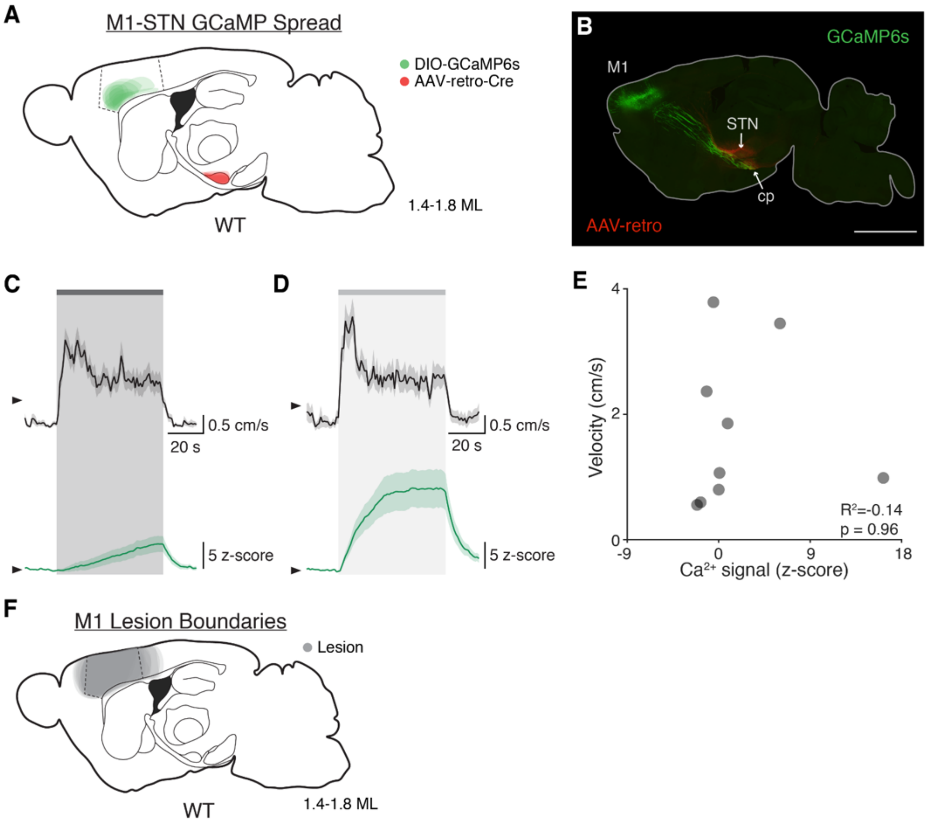


**Supplementary Figure 5. Related to Figure 4. STN DBS drives inconsistent and slow changes in hyperdirect M1 activity**. **(A)** Sagittal schematic showing estimated extent of viral AAV-retro and GCaMP6s spread in the STN and M1, respectively, of WT mice (N=9 mice). **(B)** Postmortem sagittal section example showing AAV-retro expression in the STN (red), resulting in GCaMP6s expression in M1 neurons (green) projecting to the STN and cerebral peduncle (cp). **(C-D)** Average velocity over time (black) and M1-STN GCaMP signal (green) following 60 Hz STN DBS (a, N=9 mice) or 100 Hz STN DBS (b, N=8 mice). **(E)** Scatter plot comparing hyperdirect M1 photometry signal to movement velocity during 60 Hz STN DBS (each dot represents average for one mouse, N=9 mice). **(F)** Sagittal schematic showing estimated extent of M1 lesions in WT mice (N=11 mice). Statistical significance was determined using a one-way ANOVA (see Supplementary

Table 1 for detailed statistics). Arrowhead in velocity traces and GCaMP traces corresponds to 1 cm/s and 0 z-score, respectively. Velocity and GCaMP traces show mean ± SEM.

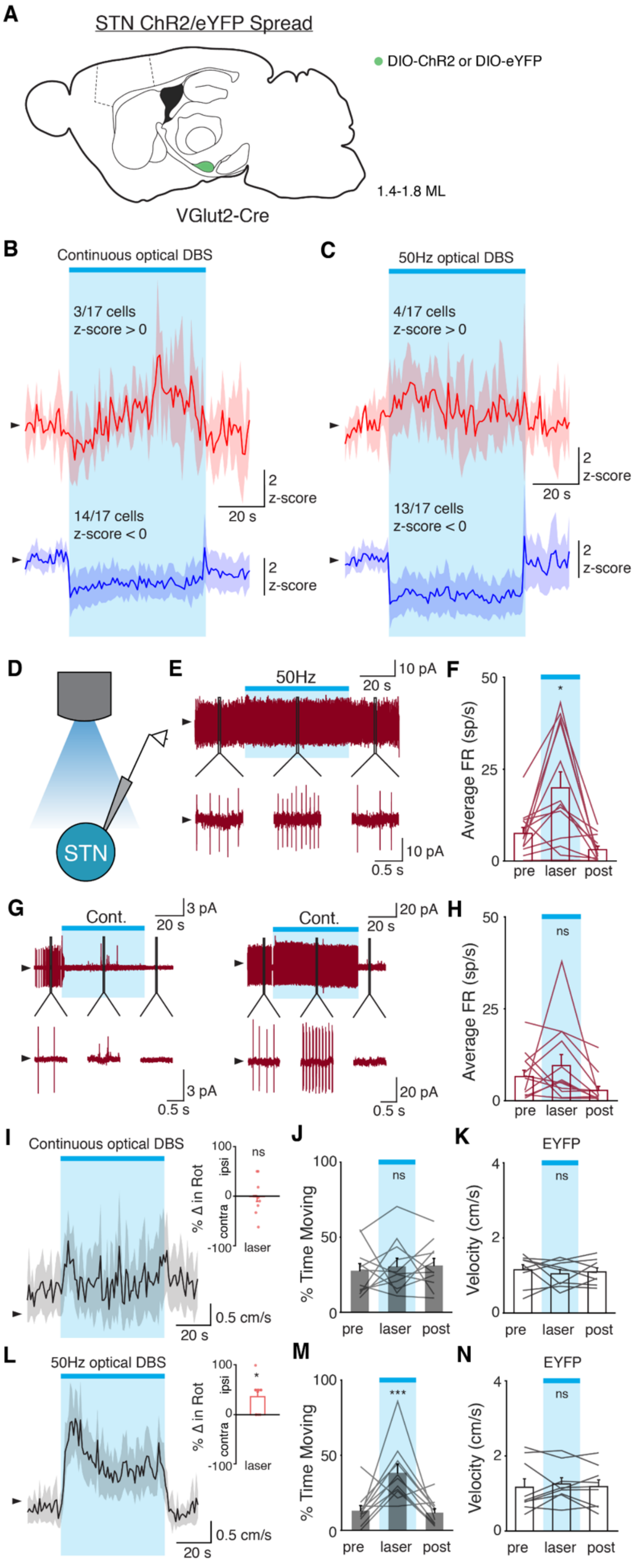

A
STN ChR2/eYFP Spread
DIO-ChR2 or DIO-eYFP
1.4-1.8 ML
VGlut2-Cre
B
Continuous optical DBS
3/17 cells
z-score > 0
14/17 cells
z-score < 0
2
z-score
20 s
C
50Hz optical DBS
4/17 cells
z-score > 0
13/17 cells
z-score < 0
D
STN
E
50Hz
10 pA
20 s
0.5 s
F
Average FR (sp/s)
pre laser post
G
Cont.
3 pA
20 pA
H
ns
I
Continuous optical DBS
% Δ in Rot
ipsi
contra
laser
0.5 cm/s
J
% Time Moving
K
EYFP
Velocity (cm/s)
L
50Hz optical DBS
M
***
N
EYFP

**Supplementary Figure 6. Related to Figure 7. *In vivo* and *ex vivo* optical STN stimulation produces distinct changes in firing rate**. **(A)** Sagittal schematic showing estimated extent of viral ChR2 or eYFP spread in the STN of VGlut2-Cre mice (N=20 mice). **(B-C)** Average *in vivo* firing rate of STN single units over time in response to continuous (B) or 50 Hz (C) optical stimulation (n=17 cells, N=3 mice), subdivided into neurons in which firing rate increased to greater than 0 z-score (top traces, red) and neurons in which firing rate decreased to less than 0 z-score (bottom traces, blue). **(D)** Recording configuration for *ex vivo* recordings of STN neurons in the cell-attached configuration. **(E)** Representative STN neuron before, during, and after 50 Hz optical stimulation. 1-second portions of the sweep are shown below. **(F)** Average firing rate before, during, and after 50 Hz optical stimulation (n=13 cells, N=3 mice). **(G)** Two STN neurons before, during, and after constant optical stimulation, demonstrating the variable response of neurons. 1-second portions of the sweep are shown below. **(H)** Average firing rate before, during, and after constant optical stimulation (n=13 cells, N=3 mice). **(I)** Average movement velocity and change in rotation bias (inset) in response to constant optical stimulation in ChR2 mice (N=11 mice). **(J)** Average percent time moving before, during, and after constant optical stimulation in ChR2 mice (N=11 mice). **(K)** Average movement velocity before, during, and after constant optical stimulation in mice injected with eYFP (N=9 mice). **(L)** Average movement velocity and change in rotation bias (inset) in response to 50 Hz optical stimulation in ChR2 mice (N=11 mice). **(M)** Average percent time moving before, during, and after 50 Hz optical stimulation in ChR2 mice (N=11 mice). **(N)** Average movement velocity before, during, and after 50 Hz optical stimulation in mice injected with eYFP (N=9 mice). Statistical significance was determined using a one-way repeated measures ANOVA with a Tukey HSD post hoc analysis applied to correct for multiple comparisons; *$P<0.05$ (only comparison between pre and laser shown, see Supplementary Table 1 for detailed statistics). Arrowhead in firing rate, cell-attached, and velocity traces corresponds to 0 z-score, 0 pA, and 1 cm/s, respectively. Firing rate traces, velocity traces, and bar plots show mean ± SEM.

### Supplementary Tables

| Figure | Part | Test | P-value (stim/las | P-value (pre vs p | P-value (stim/lase | doF (ANOVA) | F-value (ANOVA) | Cohen's d |
|---|---|---|---|---|---|---|---|---|
| 2 | D (fib) | rmANOVA | 1.19E-07 | 8.03E-01 | 1.19E-07 | 2,8 | 60.057 | 1.83 |
| 2 | D (vel) | | 7.11E-09 | 9.96E-01 | 1.09E-09 | 2,8 | 57.668 | 1.74 |
| 2 | F (fib) | | 9.54E-07 | 4.27E-01 | 4.17E-07 | 2,8 | 37.391 | 1.61 |
| 2 | F (vel) | | 1.11E-09 | 9.96E-01 | 1.09E-09 | 2,8 | 66.612 | 1.79 |
| 2 | H (fib) | | 9.04E-01 | 8.06E-01 | 8.34E-01 | 2,8 | 0.27308 | |
| 2 | H (vel) | | 1.66E-03 | 8.75E-03 | 1.02E-05 | 2,8 | 27.391 | 1.43 |
| 2 | J (spike) | | 2.89E-02 | 9.97E-01 | 5.03E-03 | 2,10 | 11.046 | 1.54 |
| 2 | J (vel) | | 3.13E-04 | 1.25E-01 | 1.77E-05 | 2,10 | 39.742 | 2.11 |
| 3 | C (fib) | | 4.14E-05 | 9.99E-01 | 4.18E-05 | 2,6 | 25.567 | 1.66 |
| 3 | C (vel) | | 1.33E-05 | 1.70E-01 | 6.55E-06 | 2,6 | 41.02 | 0.76 |
| 3 | E (fib) | | 3.32E-05 | 1.11E-01 | 9.04E-05 | 2,6 | 24.39 | 1.78 |
| 3 | E (vel) | | 4.99E-06 | 4.10E-01 | 9.82E-07 | 2,6 | 27.097 | 0.78 |
| 3 | G (fib) | | 6.68E-06 | 4.57E-03 | 1.24E-04 | 2,7 | 40.141 | 5.01 |
| 3 | G (vel) | | 1.56E-02 | 5.13E-04 | 2.25E-04 | 2,7 | 18.622 | 1.14 |
| 3 | I (spike) | | 1.21E-02 | 3.79E-01 | 9.49E-03 | 2,8 | 15.221 | 1.87 |
| 3 | I (vel) | | 1.28E-03 | 8.38E-01 | 1.62E-04 | 2,8 | 37.4 | 2.84 |
| 4 | C (fib) | | 8.37E-02 | 9.54E-02 | 1.11E-01 | 2,8 | 4.5518 | |
| 4 | C (vel) | | 1.51E-09 | 8.72E-01 | 1.18E-09 | 2,8 | 66.475 | 1.51 |
| 4 | E (fib) | | 5.88E-05 | 1.76E-02 | 6.47E-05 | 2,7 | 23.021 | 1.17 |
| 4 | E (vel) | | 4.14E-07 | 5.73E-01 | 6.95E-07 | 2,7 | 30.395 | 1.49 |
| 5 | C | | 8.24E-03 | 8.24E-01 | 5.48E-03 | 2,10 | 10.579 | 0.67 |
| 6 | D (high) | WRS | 1.48E-11 | | | | | 3.27 |
| 6 | D (low) | | 5.42E-01 | | | | | |
| 7 | C | rmANOVA | 2.46E-02 | 1.36E-01 | 6.89E-01 | 2,16 | 4.3163 | 0.10 |
| 7 | E | | 2.40E-02 | 9.97E-01 | 4.57E-02 | 2,16 | 5.5241 | 0.11 |
| 7 | F | WRS | 5.13E-01 | | | | | |
| 7 | G | | 3.88E-02 | | | | | 0.81 |
| 7 | I | rmANOVA | 2.96E-01 | 2.45E-01 | 9.87E-01 | 2,10 | 1.7004 | |
| 7 | K | | 1.27E-08 | 4.84E-01 | 1.30E-09 | 2,10 | 51.586 | 1.41 |
| S1 | B (rot) | WRS | 0.0014 (healthy vs | (PD-STN vs PD+STN) | | | | |
| S1 | B (vel) | | 0.0000823 (health | (PD-STN vs PD+STN) | | | | |
| S2 | C | FT | 3.92E-09 | 6.08E-01 | 7.41E-07 | 2 | 216.1956 (chi^2) | |
| S2 | D | | 3.53E-09 | 5.28E-01 | 1.29E-06 | 2 | 214.4129 (chi^2) | |
| S2 | E | WSR | 7.15E-01 | | | | | |
| S2 | H | FT | 5.53E-08 | 7.87E-01 | 2.01E-06 | 2 | 189.3723 (chi^2) | |
| S2 | I | | 1.46E-08 | 5.39E-01 | 4.70E-06 | 2 | 195.1556 (chi^2) | |
| S2 | J | WSR | 4.68E-01 | | | | | |
| S2 | M | rmANOVA | 9.69E-10 | 5.11E-07 | 9.56E-10 | 2,29 | 92.27 | |
| S2 | N | | 9.56E-10 | 2.02E-08 | 9.56E-10 | 2,29 | 164.19 | |
| S2 | O | WSR | 1.73E-06 | | | | | |
| S3 | D | | 3.91E-03 | | | | | |
| S3 | F (fib) | rmANOVA | 8.38E-02 | 1.18E-03 | 8.13E-01 | 2,8 | 5.6691 | |
| S3 | F (rot) | WSR | 2.89E-01 | | | | | |
| S3 | F (vel) | rmANOVA | 1.15E-03 | 1.41E-04 | 9.63E-03 | 2,8 | 22.701 | |
| S3 | G | WSR | 9.77E-04 | | | | | |
| S4 | D | | 7.81E-03 | | | | | |
| S4 | F (fib) | rmANOVA | 3.07E-01 | 2.65E-01 | 9.64E-01 | 2,7 | 1.6116 | |
| S4 | F (rot) | WSR | 5.47E-01 | | | | | |
| S4 | F (vel) | rmANOVA | 4.66E-05 | 2.04E-04 | 2.64E-02 | 2,7 | 24.425 | |
| S4 | G | WSR | 3.91E-03 | | | | | |
| S5 | E | ANOVA | 9.60E-01 | | | | 0.00336 | |
| S6 | F | rmANOVA | 3.26E-02 | 2.68E-02 | 6.86E-03 | 2,12 | 11.298 | |
| S6 | H | | 6.26E-01 | 5.81E-03 | 7.39E-02 | 2,12 | 3.6871 | |
| S6 | I | WSR | 8.10E-01 | | | | | |
| S6 | J | rmANOVA | 6.53E-01 | 4.55E-01 | 9.77E-01 | 2,10 | 0.75399 | |
| S6 | K | | 6.64E-01 | 9.31E-01 | 9.08E-01 | 2,8 | 0.31701 | |
| S6 | L | WSR | 1.60E-02 | | | | | |
| S6 | M | rmANOVA | 5.56E-09 | 7.33E-01 | 1.07E-09 | 2,10 | 52.8 | |
| S6 | N | | 8.14E-01 | 9.94E-01 | 8.21E-01 | 2,8 | 0.22566 | |

**Supplementary Table 1.** Full statistical results for the indicated figures. rmANOVA=one-way repeated measures Analysis of Variance with post-hoc Tukey multiple-comparisons test. WRS=Wilcoxon Rank Sum test (Mann-Whitney U test). FT=Friedman's test with post-hoc Tukey multiple-comparisons test. WSR=Wilcoxon Signed-Ranks test.

| **Current Amplitude (μA)** | **Frequency (Hz)** | **Pulse Width (μs)** | **High vs Low Effect** |
|---|---|---|---|
| 200 | 60 | 60 | High |
| 200 | 60 | 100 | High |
| 200 | 80 | 60 | High |
| 200 | 100 | 60 | High |
| 200 | 120 | 60 | High |
| 200 | 140 | 60 | High |
| 200 | 160 | 60 | High |
| 225 | 80 | 80 | High |
| 175 | 20 | 50 | Low |
| 200 | 10 | 120 | Low |
| 200 | 120 | 20 | Low |
| 300 | 10 | 60 | Low |
| 400 | 1 | 120 | Low |

**Supplementary Table 2. Related to Figure 6.** Stimulation parameters used in assessing differences between high and low effect stimulation.
